# Preclinical evaluation of a SARS-CoV-2 mRNA vaccine PTX-COVID19-B

**DOI:** 10.1101/2021.05.11.443286

**Authors:** Jun Liu, Patrick Budylowski, Reuben Samson, Bryan D. Griffin, Giorgi Babuadze, Bhavisha Rathod, Karen Colwill, Jumai A. Abioye, Jordan A Schwartz, Ryan Law, Lily Yip, Sang Kyun Ahn, Serena Chau, Maedeh Naghibosadat, Yuko Arita, Queenie Hu, Feng Yun Yue, Arinjay Banerjee, Karen Mossman, Samira Mubareka, Robert A. Kozak, Michael S. Pollanen, Natalia Martin Orozco, Anne-Claude Gingras, Eric G. Marcusson, Mario A. Ostrowski

**Affiliations:** Department of Medicine, University of Toronto; Toronto, ON, Canada; Institute of Medical Science, University of Toronto; Toronto, ON, Canada; Department of Molecular Genetics, University of Toronto; Toronto, ON, Canada; Lunenfeld-Tanenbaum Research Institute at Mount Sinai Hospital, Sinai Health System; Toronto, ON, Canada; Sunnybrook Research Institute; Toronto, ON, Canada; Providence Therapeutics Holdings, Inc.; Calgary, Alberta, Canada; Department of Immunology, University of Toronto; Toronto, ON, Canada; Vaccine and Infectious Disease Organization, University of Saskatchewan; Saskatoon, SK, Canada; Department of Veterinary Microbiology, University of Saskatchewan; Saskatoon, SK, Canada; Department of Biology, University of Waterloo; Waterloo, ON, Canada; Department of Medicine, McMaster University; Hamilton, ON, Canada; Department of Laboratory Medicine and Pathology, University of Toronto; Toronto, ON, Canada; Marcusson Consulting, San Francisco, CA, USA; Keenan Research Centre for Biomedical Science of St. Michael’s Hospital, Unity Health Toronto; Toronto, ON, Canada

**Author notes:** Co-first author (Jun Liu, Patrick Budylowski, Reuben Samson). Co-senior corresponding author (Eric G. Marcusson, Mario A. Ostrowski). Correspondence should be addressed to: Jun Liu or Eric G. Marcusson or Mario Ostrowski. Jun Liu and Mario Ostrowski are at Room 6368, Medical Sciences Building, 1 King’s College Circle, Toronto, ON M5S1A8, Canada. or. Eric G. Marcusson is at Providence Therapeutics Holdings, Inc. 335 25 Street SE, Calgary, AB, Canada T2A 7H8.

## Abstract

Safe and effective vaccines are needed to end the COVID-19 pandemic caused by SARS-CoV-2. Here we report the preclinical development of a lipid nanoparticle (LNP) formulated SARS-CoV-2 mRNA vaccine, PTX-COVID19-B. PTX-COVID19-B was chosen among three candidates after the initial mouse vaccination results showed that it elicited the strongest neutralizing antibody response against SARS-CoV-2. Further tests in mice and hamsters indicated that PTX-COVID19-B induced robust humoral and cellular immune responses and completely protected the vaccinated animals from SARS-CoV-2 infection in the lung. Studies in hamsters also showed that PTX-COVID19-B protected the upper respiratory tract from SARS-CoV-2 infection. Mouse immune sera elicited by PTX-COVID19-B vaccination were able to neutralize SARS-CoV-2 variants of concern (VOCs), including the B.1.1.7, B.1.351 and P.1 lineages. No adverse effects were induced by PTX-COVID19-B in both mice and hamsters. These preclinical results indicate that PTX-COVID19-B is safe and effective. Based on these results, PTX-COVID19-B was authorized by Health Canada to enter clinical trials in December 2020 with a phase 1 clinical trial ongoing (ClinicalTrials.gov number: NCT04765436).

**One Sentence Summary:** PTX-COVID19-B is a SARS-CoV-2 mRNA vaccine that is highly immunogenic, safe, and effective in preventing SARS-CoV-2 infection in mice and hamsters and is currently being evaluated in human clinical trials.

## INTRODUCTION

COVID-19 (Coronavirus Disease 2019) caused by SARS-CoV-2 (severe acute respiratory syndrome coronavirus 2) is one of the most severe health crises in human history. Since it was first reported in December 2019, more than 155 million COVID-19 cases and 3.2 million deaths have been documented and the pandemic is still spreading (*1*). Public health measures, such as social distancing, mask wearing, contact tracing, quarantine, and national lockdowns have only partially stymied the pandemic. Some treatment regimens were shown to suppress SARS-CoV-2 replication, and/or reduce the number of severe COVID-19 cases and deaths (*2–6*). Despite these advances in prevention and treatment, safe and effective SARS-CoV-2 vaccines are ultimately needed for sustainable control of the pandemic and a return to normalcy.

With unprecedented speed, hundreds of SARS-CoV-2 vaccine candidates have been designed and produced, with 280 tested in animals since the beginning of the pandemic (*7, 8*). Among them, 13 have been approved for emergency use in humans, and dozens, including the PTX-COVID19-B reported here, are at various stages of clinical trials (*7*). Given the current world population requiring vaccination, the variable conditions of public health infrastructure in different countries, and the rapid emergence of COVID-19 variants of concern (VOCs) that may escape vaccine-induced immune responses (*9–12*), continued and concerted global efforts in SARS-CoV-2 vaccine research, development, manufacturing and deployment are required to end the COVID-19 pandemic (*13*).

SARS-CoV-2 is an enveloped positive-sense RNA virus that uses the spike protein (S) on its surface to bind the angiotensin-converting enzyme 2 (ACE2) on host cells for entry to initiate replication (*14–19*). The S protein has two subunits: S1 and S2. S1 is responsible for binding to ACE2 through its receptor-binding domain (RBD). Once bound, S1 is shed from the envelope, exposing S2, which is then inserted into the host cell membrane to mediate fusion of virus envelope and cell membrane to release the viral genetic material into the host cells for replication. In contrast to SARS-CoV and other group 2B coronaviruses, SARS-CoV-2 has a furin cleavage site between the S1 and S2 subunit, which promotes infection of cells expressing the transmembrane serine protease 2 (TMPRSS2) on their surface, e.g. human respiratory tract epithelial cells (*14, 20–22*). The S protein is also the main target of host generated neutralizing antibodies (nAb) that can inhibit SARS-CoV-2 infection, e.g. by blocking its binding to ACE2 (*23–30*). Thus, most of the current SARS-CoV-2 vaccines use S protein as the immunogen.

mRNA-based vaccines are attractive platforms for prophylactic SARS-CoV-2 vaccine candidates due to their unique advantages, including rapid large-scale production, strong immunogenicity in both humoral and cellular immunity, and ease of adaptation to tackle the emerging VOCs (*31, 32*). Two SARS-CoV-2 mRNA vaccines were the earliest to enter phase 3 clinical trials, showing both high efficacy and safety, and were the first to be approved for emergency use in humans (*33, 34*). Here, we report the preclinical results of another SARS-CoV-2 mRNA vaccine, PTX-COVID19-B. We found that PTX-COVID19-B elicited potent humoral and cellular immune responses in mice, and protected both mice and hamsters from SARS-CoV-2 challenges. Based on these results, PTX-COVID19-B was authorized by Health Canada to enter clinical trials with a phase 1 clinical trial underway (ClinicalTrials.gov number: NCT04765436).

## RESULTS

### SARS-CoV-2 mRNA vaccine candidates

We first designed 3 SARS-CoV-2 mRNA vaccine candidates and compared their immunogenicity in mice: an RBD construct (amino acids 319-541), a full-length S construct (amino acids 1-1273), and an S_furinmut_ construct in which NSPRRA (amino acids 679-684) in the full-length S were replaced with IL to remove the furin cleavage site between S1 and S2 (Fig. 1A). The coding sequences of all 3 constructs were based on the S protein from the SARS-CoV-2 Wuhan-Hu-1 isolate (GenBank accession number: MN908947.3) except for a D614G substitution in the S and S_furinmut_ constructs to match this amino acid location to that of the current dominant circulating strains (*18, 35*). The RBD construct was included for testing since it is the main target of neutralizing antibodies (*23, 24, 26-28, 30*). The S_furinmut_ construct was made as it has been shown that removing the furin cleavage sites in some viral envelope proteins can enhance their expression and stability, especially when their ectodomains are expressed as soluble proteins (*36–39*). Expression of the protein encoded by these mRNA constructs on the surface of transfected HEK293T cells was detected by flow cytometry, and in the supernatant by ELISA, using an RBD-specific neutralizing mAb COV2-2165 (*30*) (Fig. S1A and S1B). As expected, both the S and S_furinmut_ proteins encoded by the mRNA constructs were expressed on the cell surface. The expressed RBD construct was detected only in the supernatant, consistent with its expression in soluble form. Some S protein could also be detected in the supernatant of the S mRNA-transfected cells, possibly due to the furin-cleavage of the membrane-bound S protein.

**Figure 1.**
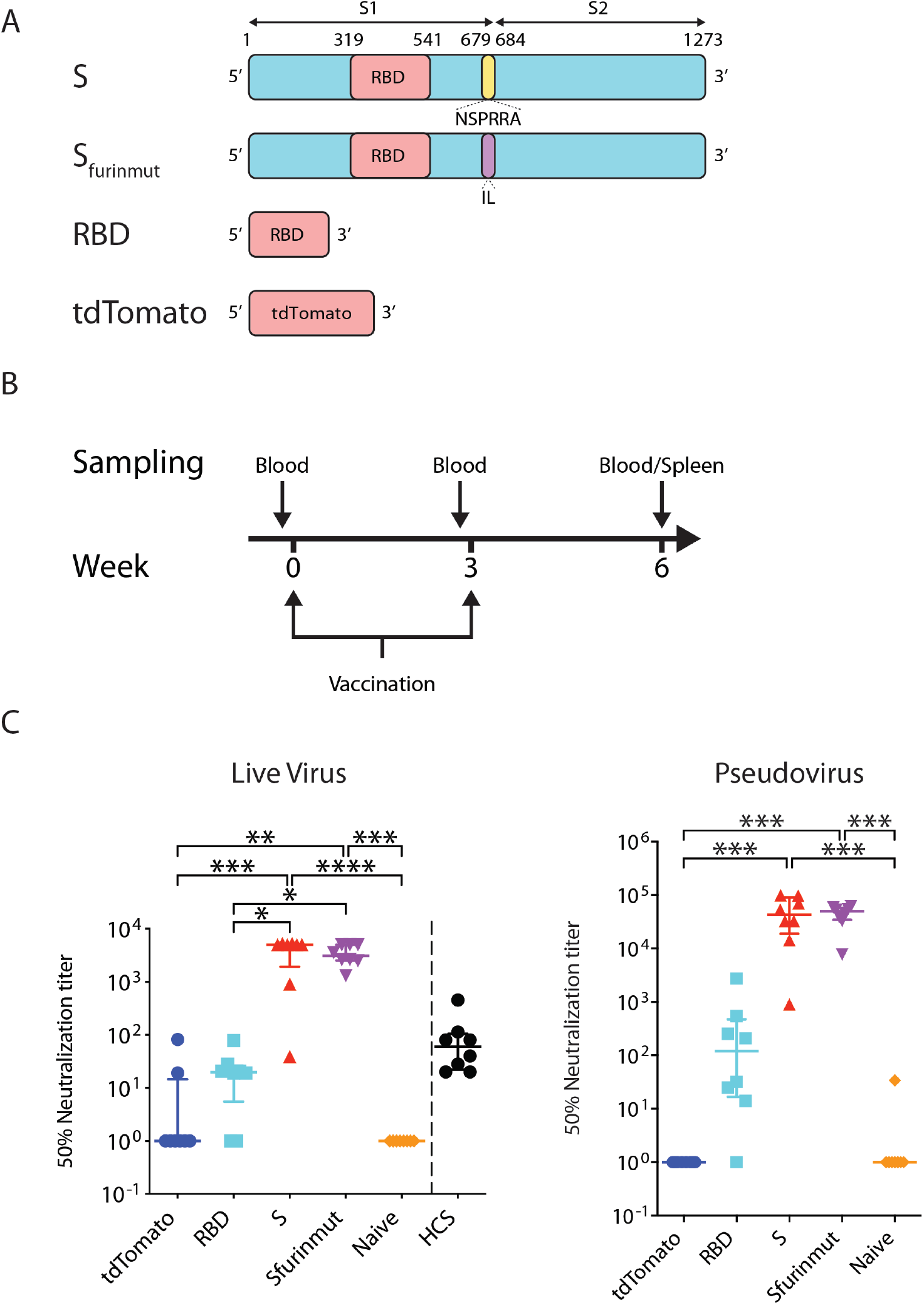
SARS-CoV-2 mRNA vaccine candidates elicits SARS-CoV-2 neutralizing antibodies in mice. **(A)** Schematic representation of the mRNA vaccine constructs. S: SARS-CoV-2 Spike mRNA (amino acids 1-1273). S_furinmut_: SARS-CoV-2 Spike mRNA (amino acids 1-1273) in which the furin cleavage site was removed by replacing NSPRRA (amino acids 679-684) with IL. RBD: SARS-CoV-2 RBD (receptor binding domain) mRNA (amino acids 319-541). tdTomato: control mRNA encoding tdTomato. S1: S1 subunit of SARS-CoV-2 Spike (amino acids 1-685). S2: S2 subunit of SARS-CoV-2 Spike (amino acids 686-1273). **(B)** Mice vaccination regimen. Six-to 8-week old mice were vaccinated twice with a 3-week interval. One day before each vaccination, peripheral blood was collected from the mice. Three weeks after the second vaccination, mice were humanly euthanized and blood and spleens were collected from the mice. **(C)** C57BL/6 mice (n=8 per group) were vaccinated with 20µg of mRNA vaccine candidates (S, S_furinmut_, RBD) or control mRNA tdTomato or DPBS for naïve control mice. Three weeks after the second vaccination, blood was collected to test neutralization of SARS-CoV-2 authentic virus or pseudovirus by the sera. For comparison, convalescent sera from 8 SARS-CoV-2 infected human subjects (HCS in the graph) were also tested for neutralization of SARS-CoV-2 authentic virus. Each symbol represents one mouse or person. Samples that did not neutralize viruses at the lowest dilution (1:20 for real virus, 1:40 for pseudovirus) are designated a 50% neutralization titer of 1. For each group, the long horizontal line indicates the median and the short lines below and above the median indicate the 25% and 75% percentile. *: *P*<0.05, **: *P*<0.01, ***: *P*<0.001, ****: *P*<0.0001 as determined by one way ANOVA (Kruskal-Wallis test) followed by Dunn’s multiple comparison test.

To compare the immunogenicity of the 3 mRNA constructs, female C57BL/6 mice were vaccinated twice, 3 weeks apart, with 20 µg of each of the constructs formulated in lipid nanoparticle (LNP) (Fig. 1B). Control mice received either 20 µg of an mRNA encoding tdTomato in the same LNP, or the same volume of Dulbecco’s phosphate-buffered saline (DPBS). Blood was collected 3 weeks post boost vaccination, and the presence of neutralizing antibodies (nAb) in the sera was measured by a micro-neutralization assay using a SARS-CoV-2 virus isolated from a SARS-CoV-2 patient (SARS-CoV-2-SB2-P3 PB Clone 1 (*40*)) (Fig. 1C). We found that the full-length S mRNA candidate elicited the highest nAb levels in the sera of vaccinated mice, followed closely by the S_furinmut_ mRNA candidate. Median nAb ID_50_ titer was 4991 (interquartile range (IQR) 1927-5188) and 3085 (IQR 2528-4991) for the S and S_furinmut_ mRNA, respectively. The RBD mRNA candidate induced low nAb levels (median 19.8, IQR 5.5-26.0). No nAb was detected in the sera from control mice receiving DPBS, and low levels of nAbs were detected in the sera of 2 out of 8 control mice receiving the tdTomato mRNA (nAb ID_50_ titer was 19 and 82, respectively). Of note, the median serum nAb titer of the S and S_furinmut_ mRNA-vaccinated mice was 83.1-fold and 51.4-fold higher than that of 8 COVID-19 convalescent patients (median nAb ID_50_ titer 60, IQR 22.0-104.8, 2 patients in each category of severe, moderate, mild, and asymptomatic SARS-CoV-2 infections), respectively (Fig. 1C). Similar results were obtained when the serum nAb was measured by a lentivirus-based SARS-CoV-2 pseudovirus neutralization assay, though the nominal nAb ID_50_ titers from the pseudovirus assay were usually higher than those from the micro-neutralization assay (Fig. 1C). Based on these results, the full-length S mRNA construct, hereafter named PTX-COVID19-B, was chosen for further testing and moved into the next stages of development.

### Humoral immune responses elicited by PTX-COVID19-B vaccination

To further evaluate the immunogenicity of PTX-COVID19-B, female C57BL/6 mice were vaccinated twice, 3 weeks apart, with 1 or 10 µg doses of PTX-COVID19-B or, as control, 10 µg of LNP formulated tdTomato mRNA. Three weeks after the boost vaccination, blood and spleens were collected from the mice to measure humoral and cellular immune responses. We first used an ELISA assay to measure S-specific binding antibodies in the sera of the mice. As shown in Fig. 2A, both 1 and 10 µg doses of PTX-COVID19-B elicited very strong S-specific total IgG, IgG1, IgG2b and IgG2c responses (median EC_50_ titers for 1 and 10 µg PTX-COVID19-B are, respectively: 1.5×10^4^ (IQR 8.1×10^3^ −2.2×10^4^), 1.1×10^5^ (IQR 7.3×10^4^ −1.5×10^5^) for total IgG; 8.3×10^3^ (IQR 3.9×10^3^ −1.5×10^4^), 1.7×10^4^ (IQR 1.1×10^4^ −2.9×10^4^) for IgG1; 5.2×10^3^ (IQR 3.0×10^3^ −6.7×10^3^), 5.9×10^4^ (IQR 4.4×10^4^ −6.3×10^4^) for IgG2b; 2.2×10^4^ (IQR 1.3×10^4^ −7.6×10^4^), 1.6×10^6^ (IQR 1.1×10^6^ −3.6×10^6^) for IgG2c). The 10 µg dose of PTX-COVID19-B usually induced higher S-specific binding antibodies than the 1 µg dose. The preponderance of the Th1 antibody (IgG2b and IgG2c) over the Th2 antibody (IgG1) also indicated that PTX-COVID19-B induced a Th1-biased antibody response. Very low levels of anti-S antibodies were detected in the sera of control mice receiving the tdTomato mRNA.

**Figure 2.**
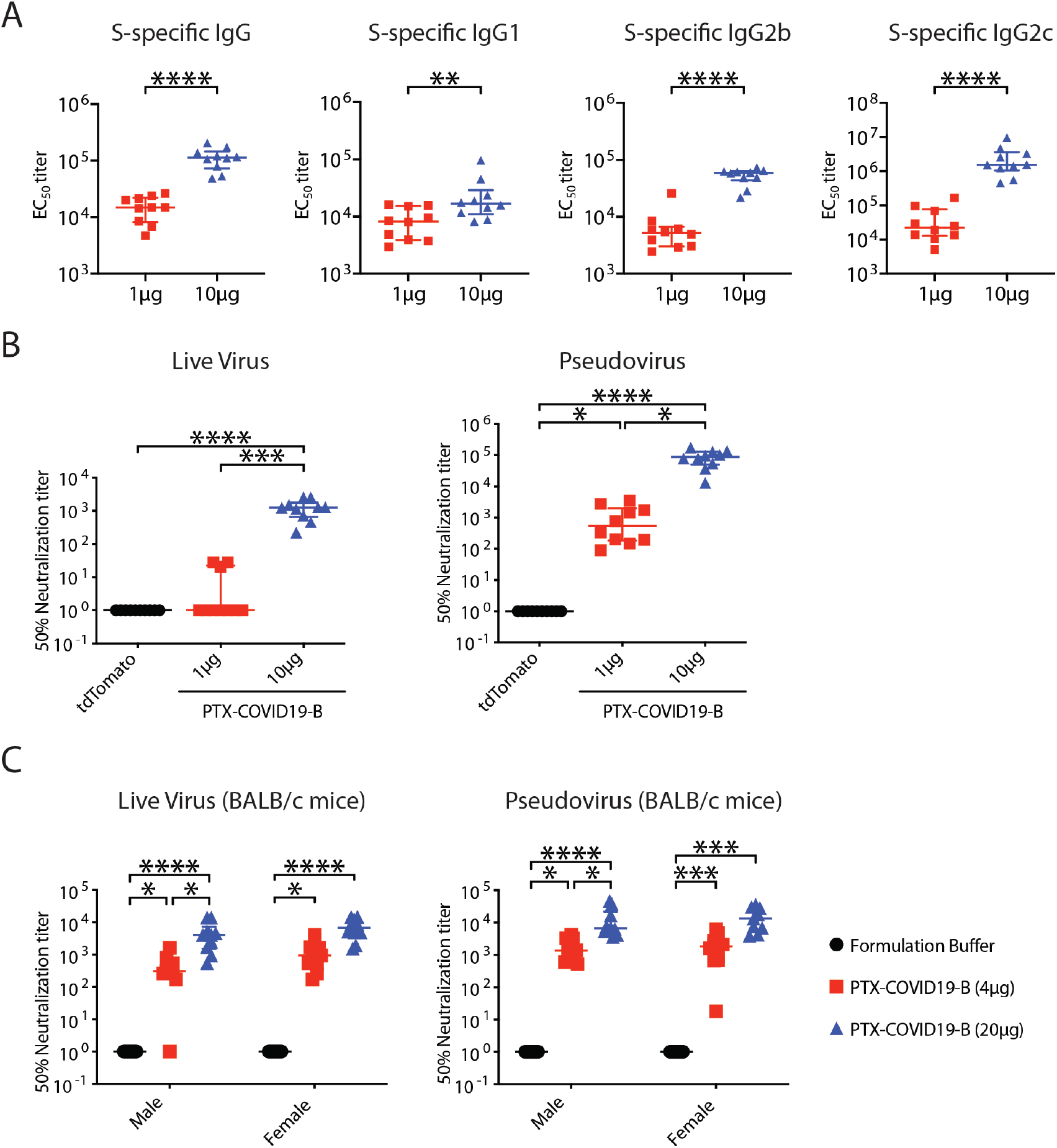
PTX-COVID19-B elicits potent humoral immune responses in mice. **(A)** and **(B)** Female C57BL/6 mice were vaccinated with 1µg or 10 µg of PTX-COVID19-B or 10 µg of control tdTomato mRNA. Three weeks after the second vaccination, blood was collected to detect (A) S-specific binding antibodies in the mouse sera as measured by ELISA and (B) neutralization of SARS-CoV-2 authentic virus or pseudovirus by the mouse sera. Shown in (A) are EC_50_ titers. N=10 for each of the PTX-COVID19-B group. Shown in (B) are 50% neutralization titers (ID_50_). N=10 per group. **(C)** Six-to 8-week old male and female BALB/c mice were vaccinated with 4 µg or 20 µg doses of PTX-COVID19-B or formulation buffer as control. Three weeks after the second vaccination, blood was collected to detect serum neutralization of SARS-CoV-2 authentic virus or pseudovirus by the mouse sera. Shown are 50% neutralization titers (ID_50_). N=10 per group except n=9 for the female 20 µg dosed PTX-COVID19-B group in the pseudovirus assay. In (B) and (C) samples that did not neutralize viruses at the lowest dilution (1:20 for real virus, 1:40 for pseudovirus) are designated a 50% neutralization titer of 1. Each symbol represents one mouse. For each group, the long horizontal line indicates the median and the short lines below and above the median indicate the 25% and 75% percentile. *: *P*<0.05, **: *P*<0.01, ***: *P*<0.001, ****: *P*<0.0001 as determined by one way ANOVA (Kruskal-Wallis test) followed by Dunn’s multiple comparison test.

We then measured nAb against SARS-CoV-2 in these C57BL/6 mouse sera. Results of SARS-CoV-2 authentic virus micro-neutralization assay showed that the 10 µg dose of PTX-COVID19-B elicited high nAb levels (median nAb ID_50_ titer was 1259, IQR 652.7-1770), which was 21.0-fold higher than that of the 8 COVID-19 convalescent patients (Fig. 2B and 1C). Low levels of nAb were induced by the 1 µg dose of PTX-COVID19-B, which, for the majority of mice, could only be detected by the pseudovirus assay, which is more sensitive (Fig. 2B). No detectable nAb was present in the sera of the mice receiving tdTomato mRNA by either assay.

To further verify the ability of PTX-COVID19-B in inducing a nAb response against SARS-CoV-2 virus, we vaccinated a different strain of mice, BALB/c, and included both sexes in the vaccination, using the same vaccination regimen as described above. Three weeks after the boost vaccination, sera were collected and nAb levels were measured. As shown in Fig. 2C, both 4 µg and 20 µg doses of PTX-COVID19-B elicited potent nAb responses in both male and female BALB/c mice. The 20 µg dose of PTX-COVID19-B induced higher nAb titers than 4 µg, although this only reached statistical significance in male mice. No detectable nAb was detected in the sera of the control mice receiving formulation buffer.

### Neutralization of VOCs by PTX-COVID19-B elicited immune sera

VOCs evade neutralization by sera from SARS-CoV-2 vaccinees, raising concerns about the efficacy of current SARS-CoV-2 vaccines. Using the pseudovirus assay (here, lentivirus particles pseudotyped to harbor the same mutations in the S protein that are found in circulating VOCs), we measured neutralization of VOCs by immune sera from PTX-COVID19-B vaccinated C57BL/6 mice. These VOCs include the B.1.1.7 lineage first detected in UK (*41*), the B.1.351 lineage in South Africa (*42*), and the P.1 lineage in Brazil (*43*) (Fig. 3A). As shown in Fig. 3B and 3C, compared to the Wuhan-Hu-1 pseudotyped lentivirus, B.1.1.7 pseudovirus was slightly resistant to neutralization by the mouse immune sera but this difference did not reach statistical significance. However, B.1.351 and P.1 pseudoviruses significantly reduced the neutralizing potency of the immune sera. For example, median serum nAb ID_50_ titer of the mice receiving a 10µg dose of PTX-COVID19-B was decreased by 25.0-fold and 11.1-fold against B.1.351 and P.1 pseudoviruses respectively, compared to Wuhan-Hu-1 pseudovirus. It should be noted that the nominal serum nAb ID_50_ titers of the immune sera from 10µg PTX-COVID19-B vaccinated mice against these VOCs are still very high, with median titers ranging from 3.2×10^3^ to 6.4×10^4^ for the different VOCs tested.

**Figure 3.**
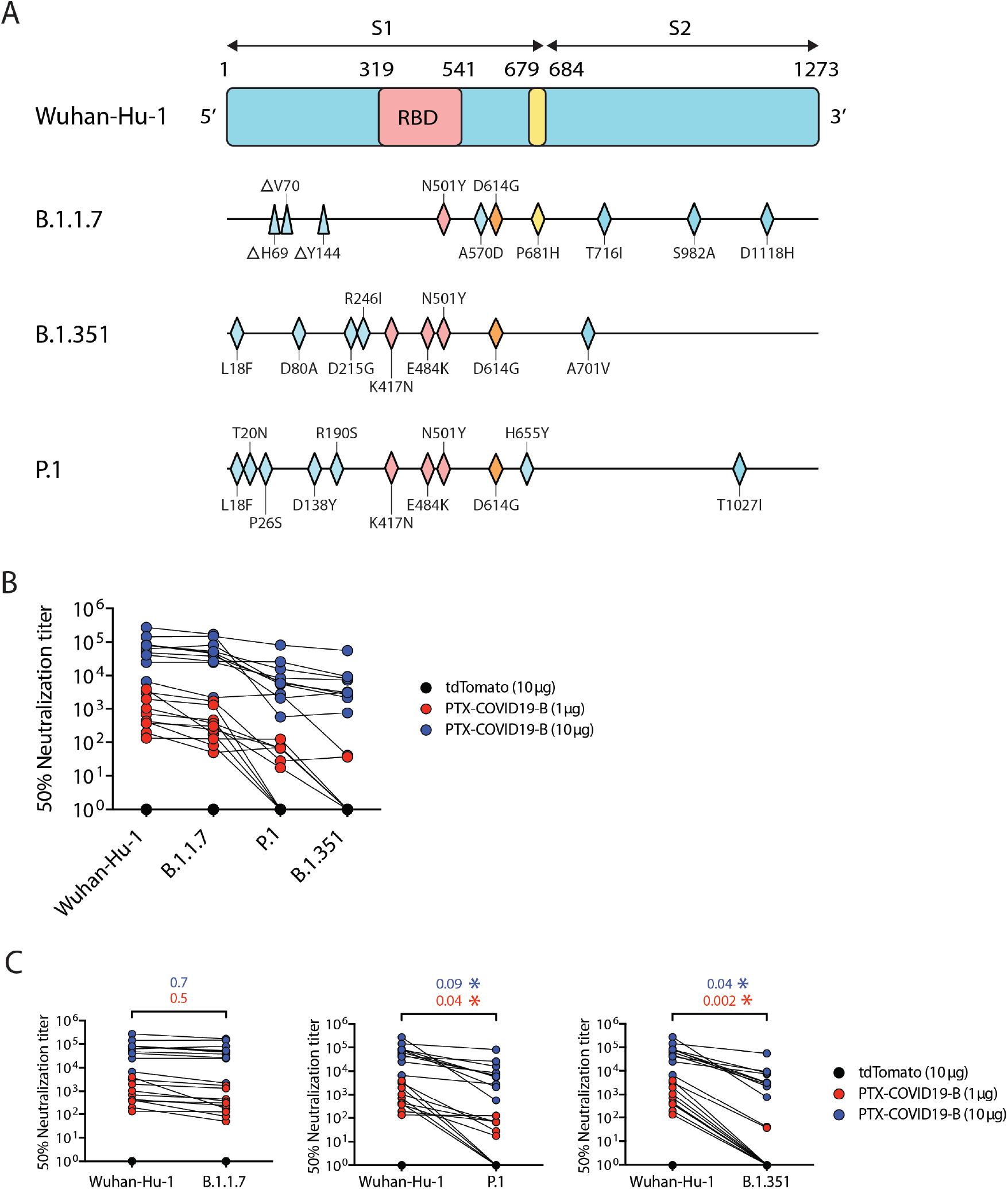
Neutralization of VOCs by immune sera from PTX-COVID19-B vaccinated mice. **(A)** Schematic representation of S proteins from SARS-CoV-2 VOCs, highlighting mutated amino acids compared to the ancestral Wuhan-Hu-1 isolate (triangles denoting deletions and rhombuses denoting replacements). **(B)** and **(C)** Neutralization of pseudoviruses bearing S protein from VOCs and Wuhan-Hu-1 isolate. C57BL/6 mice immune sera shown in Fig. 2 A and 2B were used in the neutralization. Shown in (B) are 50% neutralization titers across all pseudoviruses. Shown in (C) are pair-wise comparisons of the 50% neutralization titers between VOCs and the Wuhan-Hu-1 isolate. The numbers above the brackets in (C) are the ratios of the median 50% neutralization titers against the VOCs to the titers against Wuhan-Hu-1 isolate (blue: 10 µg PTX-COVID19-B group, red: 1 µg PTX-COVID19-B group). Each symbol represents one mouse. *: *P*<0.05 as determined by two-tailed paired t test.

### Cellular immune responses elicited by PTX-COVID19-B vaccination

C57BL/6 mice vaccinated with 1 µg and 10 µg of PTX-COVID19-B were humanly euthanized 21 days after the boost vaccination and splenocytes were stimulated with an S peptide pool (315 15-mer peptides with 11-amino-acid overlaps encompassing the entire S protein) to measure IFN-γ producing cells by ELISPOT (Fig. 4A). The 1 µg and 10 µg PTX-COVID19-B vaccinated mice had 2356±369.7 and 2810±280.9 (mean±SEM) IFN-γ spot-forming units per million splenocytes respectively, suggesting a strong Th1 response. Moreover, when splenocytes from both sexes of BALB/c mice immunized with 4 µg and 20 µg of PTX-COVD19-B were evaluated via IFN-γ and IL-4 ELISPOT, several hundreds of IFN-γ spot-forming units per million splenocytes on average were detected in immunized mice while very few IL-4 spot-forming units above the background were detected (Fig. 4B). This indicates a strong Th1 response driven by the vaccination even in a mouse strain (BALB/c) with a tendency for Th2 responses.

**Figure 4.**
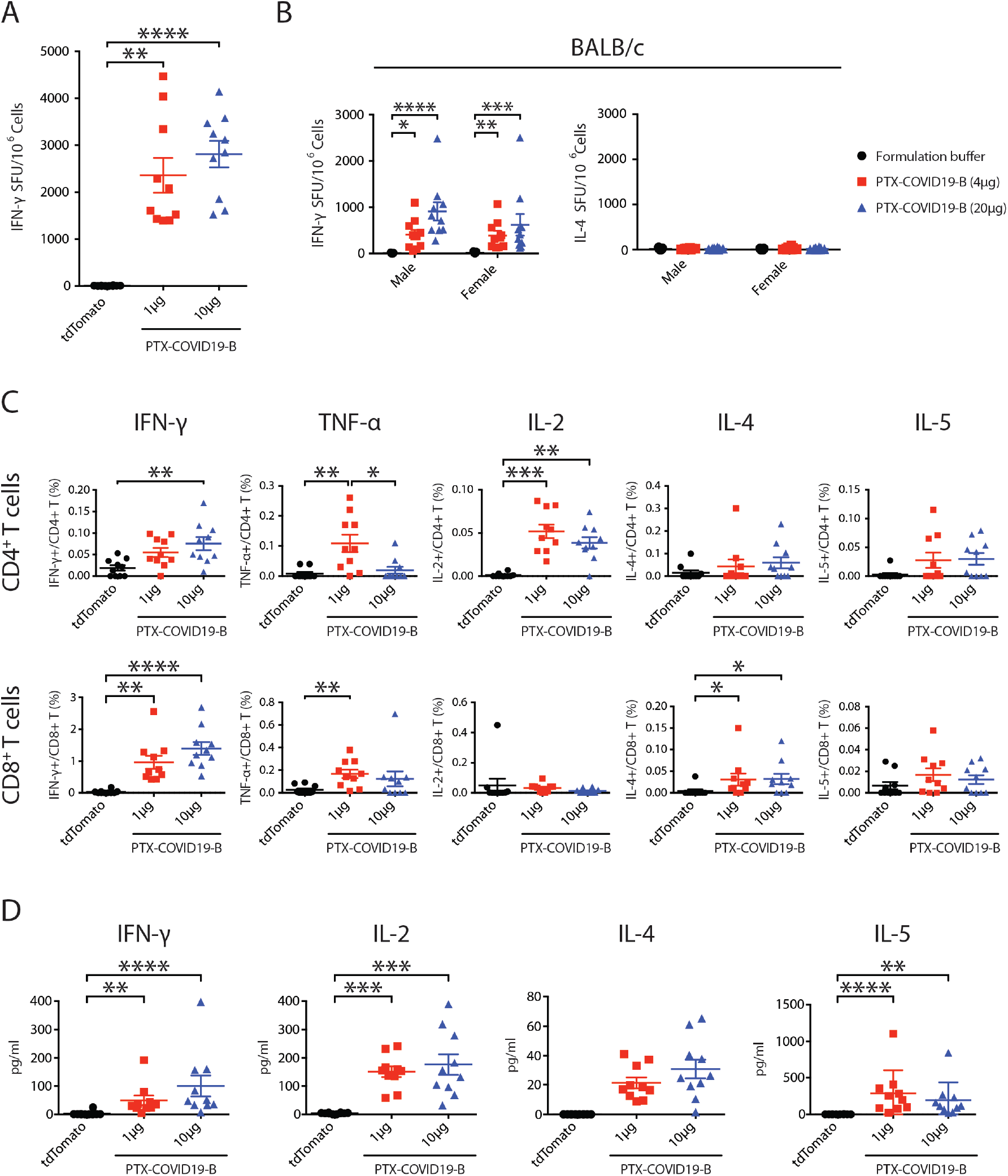
PTX-COVID19-B elicits robust cellular immune responses in mice. **(A), (C)** and **(D)** Female C57BL/6 mice were vaccinated with 1µg or 10 µg doses of PTX-COVID19-B or 10 µg of tdTomato mRNA. **(B)** Male and female BALB/C mice were vaccinated with 4 µg or 20 µg doses of PTX-COVID19-B or formulation buffer as control. Three weeks after the second vaccination, spleens were collected and the splenocytes were stimulated with SARS-CoV-2 S peptide pool to detect cytokine production as measured by ELISPOT shown in (A) and (B), intracellular cytokine staining and flow cytometry shown in (C), and a multiplex immunoassay shown in (D). Shown in (A) are the numbers of IFN-γ spot forming units (SFU) per million splenocytes (n=10 per group), (B) are the numbers of IFN-γ and IL-4 spot forming units (SFU) per million splenocytes (n=9 for each of the formulation groups, n=10 for each of the PTX-COVID19-B groups), (C) percentage of cytokine-producing cells in CD4^+^ or CD8^+^ T cells (n=10 per group), and (D) quantity of the cytokines in the supernatants of the stimulated splenocytes (n=10 per group). Each symbol represents one mouse. For each group, the long horizontal line indicates the mean and the short lines below and above the mean indicate the SEM. *: *P*<0.05, **: *P*<0.01, ***: *P*<0.001, ****: *P*<0.0001 as determined by one way ANOVA (Kruskal-Wallis test) followed by Dunn’s multiple comparison test.

Additionally, cytokine producing CD4^+^ and CD8^+^ T cells in splenocytes of C57BL/6 mice immunized with 1 and 10 µg of vaccine were analyzed by flow cytometry following overnight S peptide pool stimulation and intracellular cytokine staining (Fig. 4C). CD4^+^ T cells had increased percentages of IFN-γ, TNF-α and IL-2 producing cells, and very low percentages of IL-4 and IL-5 producing cells, indicating a strong induction of a Th1 response. Interestingly, CD8^+^ T cells showed a high number of IFN-γ producing cells, which was higher in percentage than that of CD4^+^ T cells (Fig. 4C and Fig. S2). These results contrast the T cell responses identified in COVID-19 patients, where CD4^+^ T cell responses against SARS-CoV-2 outweigh CD8^+^ T cells (*44–48*).

Furthermore, cytokines were measured in the supernatants of S peptide pool stimulated splenocytes from C57BL/6 vaccinated mice by a multiplex immunoassay (Fig. 4D), and the results confirmed a strong production of IFN-γ and IL-2 that correlates with the flow cytometry and ELISPOT data. Collectively, these results indicate that PTX-COVID19-B vaccination induced robust Th1-biased CD4^+^ and CD8^+^ T cell responses.

### PTX-COVID19-B protecting mice from SARS-CoV-2 challenge

Since wild-type mice are not susceptible to ancestral SARS-CoV-2 infection, we utilized an AAV6 (adeno-associated virus type 6)-hACE2 mouse model to test if PTX-COVID19-B can protect mice from SARS-CoV-2 infection. A similar mouse model using AAV type 9 to transduce hACE2 into mice was reported to support SARS-CoV-2 replication in mouse lungs (*49*). In our model, mice were first transduced with AAV6-hACE2 intranasally to express hACE2 in their respiratory tracts and 9 days later were intranasally inoculated with SARS-CoV-2 (Fig. 5A and Fig. S3). As shown in Fig. S3 and consistent with the previous report, AAV6-mediated hACE2 transduction induced susceptibility to SARS-CoV-2 infection as shown by the detection of infectious SARS-CoV-2 in the lungs of AAV6-hACE2 mice but not in control mice transduced with AAV6-luciferase (Fig. S3A). Using a real-time RT-PCR assay targeting the SARS-CoV-2 envelope (E) gene, we also detected a high amount of SARS-CoV-2 genomic RNA in the lungs from both AAV6-hACE2 and AAV6-luciferase transduced mice, although the genomic RNA copy numbers were much lower in the lungs of the AAV6-luciferase transduced mice than the AAV6-hACE2 mice (Fig. S3B).

**Figure 5.**
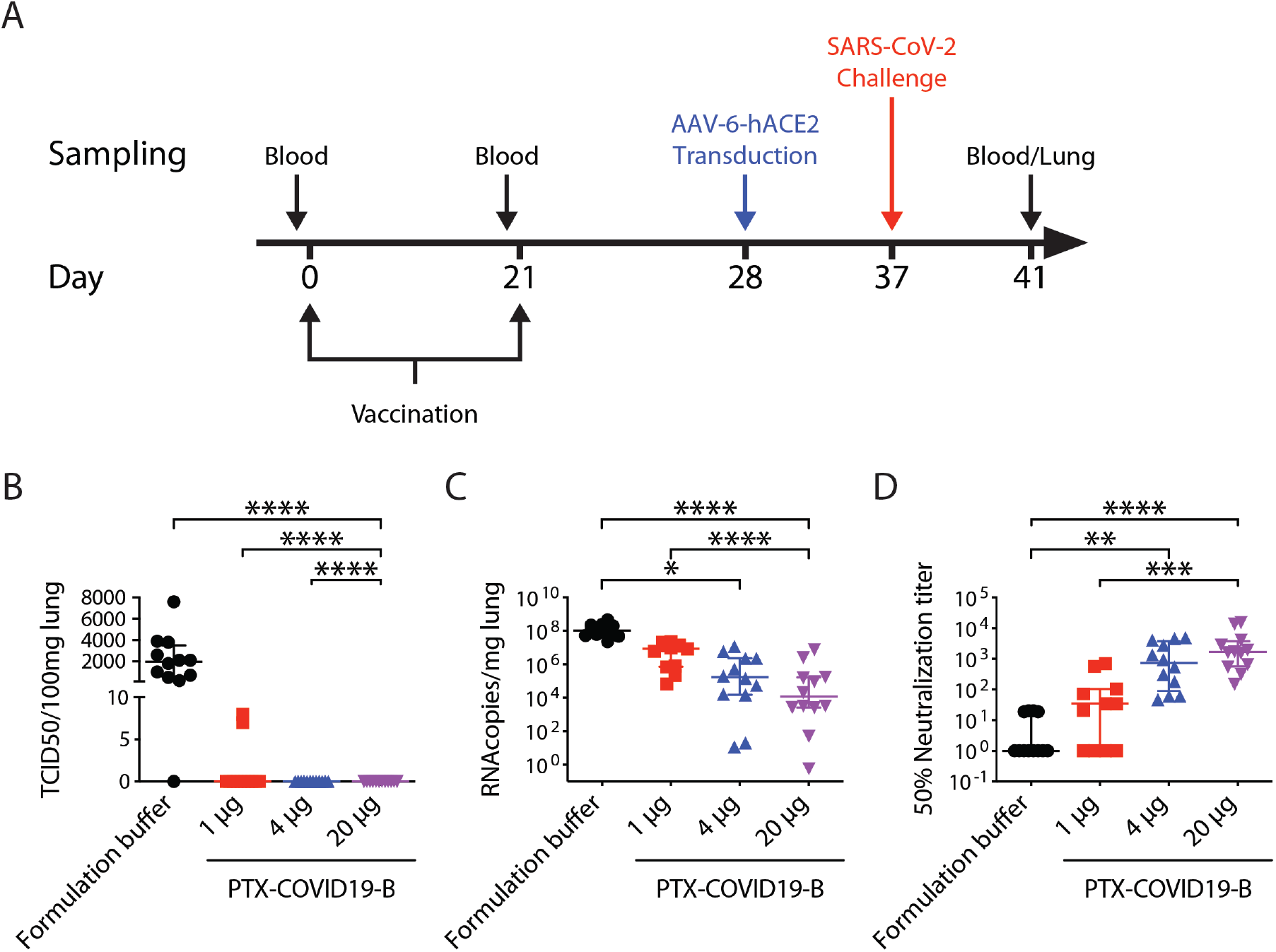
PTX-COVID19-B protects mice from SARS-CoV-2 challenge. **(A)** Mice vaccination and challenge regimen. Six-to 8-week old female C57BL/6 mice (n=10 per group) were vaccinated twice with 1µg, 4 µg or 20 µg doses of PTX-COVID19-B or formulation buffer as control. One week after the second vaccination, mice were intranasally transduced with AAV6-hACE2. Nine days after the transduction, mice were intranasally challenged with SARS-CoV-2. One day before each vaccination, blood was collected from the mice. Four days after SARS-CoV2 challenge, mice were humanly euthanized and blood and lungs were collected from the mice. **(B)** Amount of infectious SARS-CoV-2 virus and **(C)** SARS-CoV-2 RNA in the lungs of the mice. Shown in (B) are TCID_50_/100mg lung tissue (n=10 per group) and (C) RNA copies/mg lung tissue (n=10 per group). **(D)** Neutralization of SARS-CoV-2 authentic virus by the mouse sera collected 4 days after SARS-CoV-2 challenge. Shown are 50% neutralization titers (ID_50_, n=10 per group). Samples that did not neutralize viruses at the lowest dilution (1:20) are designated a 50% neutralization titer of 1. Each symbol represents one mouse. For each group, the long horizontal line indicates the median and the short lines below and above the median indicate the 25% and 75% percentile. *: *P*<0.05, **: *P*<0.01, ***: *P*<0.001, ****: *P*<0.0001 as determined by one way ANOVA (Kruskal-Wallis test) followed by Dunn’s multiple comparison test.

Having confirmed that the AAV6-hACE2 mouse model was susceptible to SARS-CoV-2 infection, we vaccinated 4 groups of C57BL/6 mice twice with 3 different doses of PTX-COVID19-B (1 µg, 4 µg and 20 µg), and as control, the formulation buffer for PTX-COVID19-B (Fig. 5). One week after the boost vaccination, all mice were transduced with AAV6-hACE2 followed by SARS-CoV-2 challenge 9 days later. Four days post challenge (4 dpi), lungs were collected and infectious SARS-CoV-2 virus in the lung tissue homogenates was quantified. Infectious SARS-CoV-2 virus was present in the lungs from 11 out of 12 control mice receiving the formulation buffer (Fig. 5B, median TCID_50_/100mg lung was 1950, IQR 550-3500). In contrast, no infectious virus was detected in the lungs from mice vaccinated with 4 µg or 20 µg doses of PTX-COVID19-B. For the mice receiving a 1 µg dose of PTX-COVID19-B, low levels of infectious virus was found in only 2 out of 12 mice (TCID_50_/100mg lung=7 and 8, respectively). SARS-CoV-2 genomic RNA could be detected in the lungs of all mice using the E-gene specific real time RT-PCR assay, but was reduced on average by 166-fold, 75-fold, and 16-fold in the mice receiving 20 µg, 4 µg, and 1 µg doses of PTX-COVID19-B, respectively, compared to the mice receiving formulation buffer (Fig. 5C). We also measured the nAb titers in the sera collected on 4 dpi and found high levels of nAb in the sera from the mice vaccinated with 20 µg and 4 µg doses of PTX-COVID19-B and moderate levels of nAb from the mice receiving a 1 µg dose of PTX-COVID19-B (Fig. 5D). Given the short time after SARS-CoV-2 challenge (4 dpi), the nAb levels in these mouse sera was most likely elicited by the vaccination, not induced or boosted by the SARS-CoV-2 infection. Of note, the serum nAb ID_50_ titers negatively correlate with the quantities of the infectious SARS-CoV-2 virus and the genomic RNA in the lungs (Fig. S4). Logistic regression modeling of the nAb ID_50_ titers and the virus TCID_50_ values indicates that a threshold nAb ID_50_ titer of 654.9 against authentic virus predicts a 95% probability of protection from productive SARS-CoV-2 infection. Taken together, these data indicate that PTX-COVID19-B completely protected mice from pulmonary infection by SARS-CoV-2, even at a low dose of 4 µg.

### PTX-COVID19-B protecting hamsters from SARS-CoV-2 challenge

Syrian hamsters are susceptible to and can transmit SARS-CoV-2 infection, mimicking some aspects of SARS-CoV-2 infection in humans (*50, 51*). We thus tested the efficacy of PTX-COVID19-B in Syrian hamsters (Fig. 6). Two groups of hamsters (n=8) were vaccinated twice with a 3-week interval with either a 20 µg dose of PTX-COVID19-B or the formulation buffer. Twenty days after boost vaccination, all hamsters were challenged intranasally with SARS-CoV-2. Body weight of the hamsters was measured 1 day before the SARS-CoV-2 challenge and then on 1, 3, 5, 7 and 8 dpi. Oral swabs were taken from the hamsters on 1, 3, 5 and 7 dpi. On 4 and 8 dpi, half (n=4) of the hamsters from each group were humanly euthanized, and nasal turbinates and lungs were collected.

**Figure 6.**
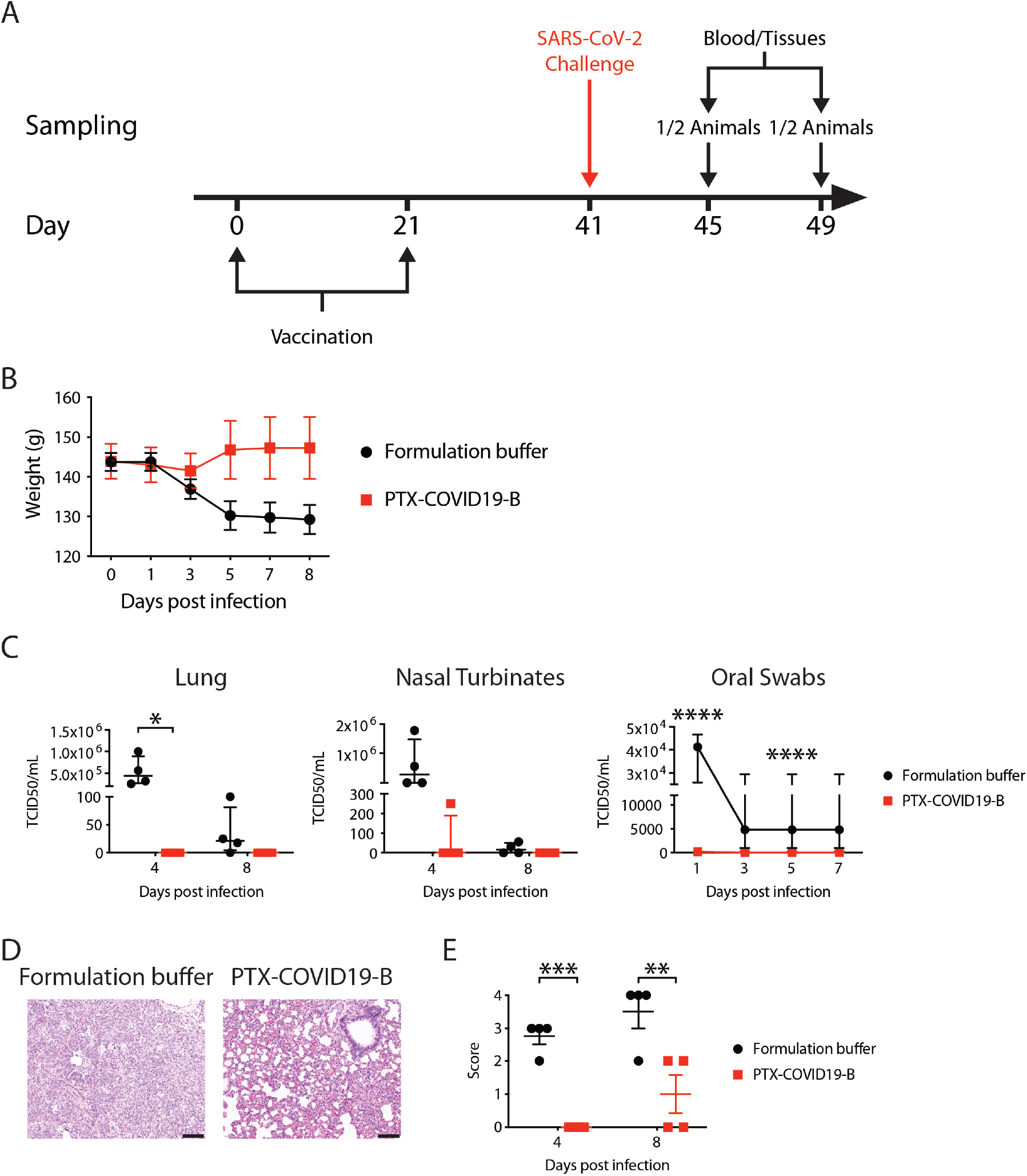
PTX-COVID19-B protects hamsters from SARS-CoV-2 challenge. **(A)** Hamster vaccination and SARS-CoV-2 challenge regimen. Six-to 10-week old male Syrian hamsters (n=8) were vaccinated with a 20 µg dose of PTX-COVID19-B or formulation buffer twice with a 3-week interval. Twenty days after the second vaccination, hamsters were challenged intranasally with SARS-CoV-2. Four days and 8 days after the challenge, half of the animals in each group (4 animals per group) were humanly euthanized and blood and tissues were collected. **(B)** Body weight of the hamsters. Symbols indicate the means, and the error bars indicate SEM for each group. N=8 per group from day 0 to day 3. N=4 from day 5 to day 8. **(C)** Amount of infectious SARS-CoV-2 virus in the hamster tissues. Shown are TCID_50_ per ml of the tissue homogenates (lung and nasal turbinates) or per ml of oral swab samples. For lungs and nasal turbinates, each symbol represents one hamster (n=4 per group per time point), and the long horizontal line indicates the median and the short lines below and above the median indicate the 25% and 75% percentile. For oral swabs, symbols indicate the medians, and the error bars indicate the 25% and 75% percentile. (n=8 per group from day 0 to day 3 and n=4 from day 5 to day 7). **(D)** Representative micrographs of the hamster lungs, showing extensive acute and mixed inflammatory cell infiltrates in bronchiole and alveoli in a control hamster receiving formulation buffer and paucity of inflammation in a vaccinated animal after challenged with SARS-CoV-2. Scale bar=100µm. **(E)** Pathology score of the hamster lungs. Shown are semiquantitative pathology scores of the hamster lungs. Each symbol represents one hamster (n=4 per group per time point). The long horizontal lines indicate the means and the short lines below and above the mean indicate the SEM. *: *P*<0.05, **: *P*<0.01, ***: *P*<0.001, ****: *P*<0.0001 as determined by two way ANOVA followed by Sidak’s multiple comparison test.

When compared to pre-challenge, the body weight of the control hamsters decreased from 3 dpi to 8 dpi, while that of the PTX-COVID19-B vaccinated hamsters decreased slightly on 3 dpi and then increased from 4 dpi to 8 dpi (Fig. 6B). We then measured the amount of infectious virus in the respiratory tracts of the hamsters (Fig. 6C). No infectious SARS-CoV-2 virus was detected in the lungs of PTX-COVID19-B vaccinated hamsters on 4 dpi and 8 dpi. In the lungs of control hamsters, a large amount of infectious virus was found on 4 dpi (median TCID_50_=4.4×10^5^, IQR 2.7×10^5^-8.9×10^5^), and low levels of infectious SARS-CoV-2 virus (median TCID_50_=21.5, IQR 4.4-81.3) were still present in 3 out of 4 animals on 8 dpi. Infectious virus was also found in the nasal turbinates of control hamsters on 4 dpi and 8 dpi, respectively (4 dpi median TCID_50_=2.8×10^5^, IQR 4.4×10^3^-1.5×10^6^; 8 dpi median TCID_50_=15.8, IQR 0-50.1). No infectious virus was detected in the nasal turbinates of PTX-COVID19-B vaccinated hamsters on 4 dpi and 8 dpi except in one animal where low level (TCID_50_=251) was detected on 4 dpi (Fig. 6C). Similar results were observed in the oral swabs, where little or no infectious SARS-CoV-2 virus was detected in the samples of PTX-COVID19-B vaccinated hamsters but high levels of the virus were detected in control hamsters from 1 dpi to 5 dpi (Fig. 6C).

Lung pathology was also examined in all hamsters, using a semiquantitative grading system to score the severity of the lung pathology (Fig. 6D, 6E, and Table S1). There was a significant difference in the lung pathology of control animals (n=4) and PTX-COVID19-B vaccinated animals (n=4), after challenge with SARS-CoV-2. The main histopathologic features after SARS-CoV-2 infection was extensive mixed inflammatory cell infiltration. The lung pathology was less extensive in the PTX-COVID19-B vaccinated animals, although there was individual variability in grades.

Taken together, these results indicate that vaccination with PTX-COVID19-B prevented productive infection of the lungs and upper respiratory tracts by SARS-CoV-2 in hamsters, and protected the animals from moderate/severe lung inflammation.

### Safety of PTX-COVID19-B in the animals

C57BL/6 mice and hamsters were checked daily during the experiments. The general status of the vaccinated animals such as appearance, feeding, and mobility was the same as the control mice.

Male and female BALB/c mice immunized with 4 µg and 20 µg doses of PTX-COVID19-B were evaluated for body weight, and injection site dermal scoring using a modified Draize scoring method (*52*) on days 23 and 43 (2 and 22 days after the second immunization, respectively). No differences in body weight were observed in immunized males compared to control mice. A small weight loss was observed in females compared to control animals in the week following the second injection of PTX-COVID19-B at 20 µg dose but by day 43, there was no significant difference in average body weight compared to control groups. A slight transient dermal erythema was observed in a small proportion (20%) of vaccinated mice, which disappeared by day 43. Therefore, PTX-COVID19-B intramuscular immunization in BALB/c mice was well tolerated and only mild transient effects were observed, which resolved by day 43.

## DISCUSSION

The results presented here indicate that the SARS-CoV-2 mRNA vaccine, PTX-COVID19-B, is safe and effective in mouse and hamster models. PTX-COVID19-B elicited robust cellular and humoral immune responses and could completely protect vaccinated mice or hamsters from productive SARS-CoV-2 infection in the lungs. Although it is difficult to make a direct comparison due to variations in experimental conditions, the immunogenicity, such as the nominal titers of vaccine-induced nAb, and the efficacy of PTX-COVID19-B, are comparable to those reported in the small animal studies of the other two mRNA vaccines approved for emergency use in humans (*32, 53*).

PTX-COVID19-B also prevented SARS-CoV-2 replication in the upper respiratory tracts of hamsters, as shown by little or no infectious virus detected in nasal turbinates or oral swabs. Suppression of virus replication in upper respiratory tracts can reduce transmission of respiratory viruses, but has been hard to achieve for respiratory virus vaccines, possibly due to poor performance of these vaccines in inducing mucosal immunity in upper respiratory tracts (*54, 55*). In this regard, some SARS-CoV-2 vaccines, including one of the two approved mRNA vaccines, were shown to be capable of suppressing the virus replication in both upper and lower respiratory tract in animal studies (*32, 53, 56, 57*). Additional experiments are needed to confirm the upper respiratory tract findings reported here, including the examination of the mucosal immune response elicited by PTX-COVID19-B and tests to see if PTX-COVID19-B can prevent SARS-CoV-2 transmission between hamsters.

PTX-COVID19-B encodes a full-length membrane-anchored S protein derived from the ancestral Wuhan-Hu-1 isolate with a D614G substitution to match the predominant circulating SARS-CoV-2 strains at this amino acid location. During the preparation of this manuscript, SARS-CoV-2 VOCs emerged and have begun dominating the circulating strains worldwide, casting a doubt on the efficacy of current SARS-CoV-2 vaccines (*41–43*). Fortunately, immune sera from human subjects receiving the other two approved mRNA vaccines have been shown to neutralize many VOCs, although usually with reduced titers, in particular for the B1.351 lineage (*9–12, 58*). Consistent with these reports, PTX-COVID19-B elicited mouse immune sera were still capable of neutralizing 3 dominant VOCs with high potency by a pseudovirus assay. However, further experiments are needed to test the efficacy of PTX-COVID19-B in protection of animals against VOCs’ infections.

The 2P mutation (K986P and V987P) was reported to stabilize the ectodomain of the S protein in the prefusion conformation (*17*), which was regarded as crucial in inducing nAb, and thus was adopted in some SARS-CoV-2 vaccines (*32, 53, 59, 60*). PTX-COVID19-B does not have the 2P mutation since we thought it likely that the membrane-anchor could stabilize the full-length S in the prefusion conformation as reported for other virus envelope proteins (*61–63*). An additional example for the use of wild-type full-length S in a SARS-CoV-2 vaccine is ChAdOx1 nCoV-19 (AZD1222), which was reported assuming the prefusion conformation on the cell surface (*64*). The equivalent levels of SARS-CoV-2 nAb titers and protection from SARS-CoV-2 infection afforded by PTX-COVID19-B, as compared to those of the other two approved mRNA vaccines using the 2P mutations in their S immunogens, suggest that the 2P mutation might not be essential for induction of protective immunity (*32, 53*). We also designed and tested another S construct, S_furinmut_, in which the furin cleavage site between S1 and S2 subunits was removed. Removing this site was presumed to stabilize the ectodomain of the S protein by keeping the S1 subunit from shedding from S, and was utilized in some SARS-CoV-2 vaccines (*17, 59, 60*). However, we did not find that the S_furinmut_ mRNA performed better in eliciting nAb responses in mice compared to the S mRNA. Additional experiments are required to test if other modifications of the S protein could enhance the strength and breadth of the immunogenicity and efficacy of PTX-COVID19-B. In contrast to the S mRNA and S_furinmut_ mRNA, the RBD mRNA performed poorly in inducing a nAb response. This is consistent with previous reports showing that RBD was weakly immunogenic, possibly due to its small size (*65, 66*).

The protection mechanisms of SARS-CoV-2 vaccines have not been fully elucidated in humans, although nAbs and T cells are assumed to be critical protection correlates (*47, 54, 67*). This is supported by animal studies where nAb responses were pivotal in protecting monkeys from SARS-CoV-2 infection, with CD8^+^ T cells also participating in protection (*68*). In addition, CD4^+^ T cell help is vital for the quantity and quality of nAb and CD8^+^ T cell responses against virus infection (*69–71*). T cells could also continue to attack the VOCs that escape the nAb response, since they recognize multiple linear epitopes, including epitopes in the conserved region of the S protein (*72*). In this regard, PTX-COVID19-B elicited both robust nAb and CD8^+^ T cell responses. PTX-COVID19-B also induced a predominant Th1 response, which is regarded as a desirable feature for respiratory virus vaccines (*73*). Additional experiments will be required to track nAb and T cell responses induced by PTX-COVID19-B, including their durability and capability to protect against VOCs.

Although a few vaccines have been approved for emergency use in humans, additional safe, effective, and easily deployable SARS-CoV-2 vaccines are needed to meet the enormous challenge for the global immunization required to end the COVID-19 pandemic. Based on the results reported here, PTX-COVID19-B has been authorized by Health Canada to enter a phase 1 clinical trial (ClinicalTrials.gov number: NCT04765436). Results of the clinical trial will determine if PTX-COVID19-B will enter into to the next phases of clinical trials and eventually be added into the vaccine arsenal in the global fight against the SARS-CoV-2.

## MATERIALS AND METHODS

### Study design

The objective of this study was to evaluate the immunogenicity, safety and efficacy of a SARS-CoV-2 mRNA vaccine in mice and hamsters. The sample size of mice was determined by power analysis assuming 60% protection efficacy. Due to the capacity limit in our animal facility, a total of 16 hamsters in two groups were used in this study. All animals were randomly assigned to different treatment groups. The performers measuring SARS-CoV-2 neutralization by mice sera and SARS-CoV-2 virus in mouse tissues and the pathologist examining animal pathology were blinded to the sample groupings.

### Ethics

All animal work was approved by the Animal Care Committees of The University of Toronto. For studies involving human samples, written informed consent was obtained from convalescent COVID-19 patients, and samples were obtained and used according to a research ethics board (REB) approved protocol (St. Michael’s Hospital REB20-044c to M.A.O.).

### Vaccine

The mRNAs used in these studies encode the full-length S (amino acids 1-1273), the S_furinmut_ that is the same as the full-length S except for the furin cleavage site NSPRRA (amino acids 679-684) changed to IL, or the RBD (amino acids 319-541). The amino acid sequences encoded by these mRNAs are the same as the Spike protein sequence from SARS-CoV-2 Wuhan-Hu-1 isolate, GenBank accession number: MN908947.3, except for a D614G substitution in S mRNA and S_furinmut_ mRNA. The mRNAs contains codon-optimized open reading frames for the S, S_furinmut_, or RBD flanked by an optimized capped 5’ UTR and an optimized 3’ UTR followed by a poly-A tail. The mRNA was produced by *in vitro* transcription of a linear plasmid template using T7 RNA polymerase. The mRNA was purified by Providence’s proprietary purification process using a series of purification steps to remove transcription enzymes, the linear DNA template and mRNA-related impurities prior to formulation. LNPs were prepared by mixing a buffered solution of mRNA with an ethanol solution of lipids (DSPC, cholesterol, PEG-lipid and ionizable lipid) following Genevant’s proprietary process (Vancouver, BC, Canada). The LNPs were concentrated by tangential flow ultrafiltration and then diafiltered against an aqueous buffer system. Following a 0.2 µm filtration process, the LNPs were subjected to quality tests including RNA concentration, encapsulation efficiency, particle size, pH and osmolality.

### mRNA transfection of HEK293T cells and detection of the expressed immunogens

S mRNA, S_furinmut_ mRNA, and RBD mRNA were transfected into HEK293T cells using Lipofectamine® MessengerMAX™ transfection reagent (Thermo Fisher Scientific, Mississauga, ON, Canada) according to the manufacturer’s protocol. Briefly, HEK293T cells cultured in DMEM-10 medium (DMEM supplemented with 10% FBS, 100U penicillin, 100µg streptomycin, and 2mM L-glutamine. DMEM was purchased from Thermo Fisher Scientific. All others were purchased from Wisent Bioproducts, St-Bruno, QC, Canada) were seeded into 6-well plates (Corning Life Sciences, Tewksbury, MA). After overnight culture, 2 µg mRNAs were diluted in 125 µl Opti-MEM (Thermo Fisher Scientific), mixed with the Lipofectamine® MessengerMAX™ transfection reagent, and then added to the cells. Twenty-four hours later, supernatant was collected from the transfected cells for an in-house sandwich ELISA to detect the expressed immunogens in the supernatant. Cells were collected for the detection of the expressed immunogens on the cell surface by flow cytometry.

For the sandwich ELISA, Immunolon 2HB flat-bottom microtiter plates (Thermo Fisher Scientific) were coated with an RBD-specific neutralizing mAb COV2-2165 (kindly provided by Dr. J.E. Crowe Jr. from Vanderbilt University Medical Center, Nashville, TN), washed, and blocked with DPBS containing 3% BSA (Sigma-Aldrich, Oakville, ON, Canada). The supernatant was then pipetted into the plates. Purified RBD protein (kindly provided by Dr. J. M. Rini, Department of Biochemistry, University of Toronto, Toronto, ON, Canada) was also added to the plates as positive control. After 2 hours incubation, the plates were washed and mouse anti-S immune serum (kindly provided by Dr. J. R. Carlyle from Department of Immunology, University of Toronto, Toronto, ON, Canada) was added to the plates. After 1-hour incubation, the plates were washed and HRP-labeled goat anti-mouse IgG (SouthernBiotech, Birmingham, AL) was added. After 1-hour incubation, SureBlue TMB microwell peroxidase substrate (KPL, Gaithersburg, MD) was added and 15 minutes later, 1N HCL was pipetted into the plates to stop the reaction. OD_450_ was then read using a microplate reader (Thermo Fisher Scientific).

For flow cytometry, cells were first stained with the RBD-specific neutralizing mAb COV2-2165, and then with an APC mouse anti-human IgG (BD Biosciences, Mississauga, ON, Canada). Stained cells were run on LSRFortessa (BD Biosciences). FlowJo (BD) was used to analyze the flow cytometry data.

### Mouse vaccination

Female C57BL/6 mice of 6-to 8-week old were vaccinated intramuscularly twice with a 3-week interval. In some experiments, both male and female BALB/c mice of 6-to 8-week old were used. Various doses of mRNA vaccines or control tdTomato mRNA in 50µl total volume were injected into the hind leg muscle for each immunization. Naïve mice received the same volume of either DPBS or the vaccine formulation buffer. Each day before vaccination, blood was collected from the mice through the saphenous vein. Three weeks after boost vaccination, mice were humanly euthanized and spleen and blood samples were collected. Serum was isolated from the blood by centrifugation at 10,000 *g* for 30 seconds at 4°C.

### S-specific immunoglobulin ELISA

ELISAs were performed as previously described with minor modifications for mouse samples(*74*). 96-well plates (Green BioResearch, Baton Rouge, LA) were coated with 200 ng/well of recombinant purified full-length spike trimer SmT1(*75*) and blocked with 3% w/v milk powder in PBS-T. Serial dilutions of mouse samples in 1% w/v milk powder in PBS-T were added to the plate (starting at 1:100 dilution with 5-fold dilutions) and incubated for 2 hours at room temperature. Wells were then washed 3 times with 200µL PBST before incubation for 1 hour with secondary antibodies (HRP-labeled Goat anti-mouse IgG1/IgG2b/IgG2c purchased from SouthernBiotech or HRP-labeled Goat anti-mouse IgG Fcγ purchased from Jackson ImmunoResearch, West Grove, PA) in 1% w/v milk powder in PBS-T. Samples were washed 3 times with PBS-T and then 1-Step Ultra TMB-ELISA Substrate Solution (Thermo Fisher Scientific) was added for 15 minutes at room temperature. The reaction was quenched with equal volume stop solution containing 0.16N sulfuric acid (Thermo Fisher Scientific). Plates were read using a spectrophotometer (BioTek Instruments Inc., Cytation 3) reading at 450nm. All sample raw OD values had blank values subtracted before analysis. OD values of each PTX-COVID19-B vaccinated mouse serum minus average of OD values of 4 tdTomato control mouse sera at the same dilution were used to calculate EC_50_ titer using the 4-parameter logistic regression analysis in GraphPad Prism 8 (GraphPad Software, La Jolla, CA).

### Serum neutralization using SARS-CoV-2 virus

A micro-neutralization assay was used to measure the neutralizing titers of the sera(*45*). Briefly, VeroE6 cells cultured in DMEM-10 were seeded into 96-well plates and cultured overnight. Sera were heat-inactivated at 56°C for 30 minutes. Serial dilutions of the sera were mixed with 100 TCID_50_ SARS-CoV-2 virus (isolate SARS-CoV-2-SB2-P3 PB Clone 1, passage 3(*40*)) in serum free DMEM, incubated at 37°C for 1 hour, and then added onto the VeroE6 cells. The cell plates were then incubated at 37°C for 1 hour, shaking every 15 minutes. Inoculums were then removed and DMEM-2 (DMEM supplemented with 2% FBS, 100U penicillin, 100µg streptomycin, and 2mM L-glutamine) was added to the cells. Cell plates were incubated at 37°C for 5 days and cytopathic effect (CPE) was checked every day. 50% neutralization titer (ID_50_) was defined as the highest dilution factor of the serum that protected 50% of the cells from CPE and calculated by using the 4-parameter logistic regression analysis in GraphPad Prism 8. The performer of the assay was blinded to the grouping of the mice.

### Serum neutralization using pseudovirus

Spike-pseudotyped lentiviral assays were performed as previously described with reagents kindly provided by Dr. J. D. Bloom (Department of Genome Sciences, University of Washington, Seattle, WA) and with minor modifications for mouse samples(*74*). Briefly, Spike-pseudotyped lentivirus particles (both wild-type Wuhan-Hu-1 and tested VOCs) were generated and used at ∼1:25 virus stock dilution (a virus dilution resulting in >1000 relative luciferase units (RLU) over control). For the neutralization assay, diluted mouse sera (1:40 from stock sera) were serially diluted (from 2.5 to 4-fold dilutions over 7 dilutions to encompass a complete neutralization curve per sample) and incubated with diluted pseudovirus at a 1:1 ratio for 1 hour at 37°C before being transferred to plated HEK293T-ACE2/TMPRSS2 cells and incubated for an additional 48 hours at 37°C and 5% CO_2_. After 48 hours, cells were lysed, and Bright-Glo luciferase reagent (Promega, Madison, WI) was added for 2 minutes before reading with a PerkinElmer Envision instrument (PerkinElmer, Waltham, MA). 50% neutralization titer (ID_50_) were calculated with nonlinear regression (log[inhibitor] versus normalized response – variable slope) using GraphPad Prism 8. The assay was performed in the same manner for all VOCs tested. The performer of the assay was blinded to the grouping of the mice.

#### ELISPOT assay

To perform IFN-γ ELISPOT for C57BL/6 mice, ELISPOT plates (Sigma-Aldrich) were coated with rat anti-mouse IFN-γ antibody (BD Bioscience) overnight. Plates were washed and blocked with RPMI-10 medium (RPMI-1640 supplemented with 10% FBS, 100U penicillin, 100µg streptomycin, and 2mM L-glutamine. All were purchased from Wisent Bioproducts) for 2 hours. Splenocytes were added into the plates, and stimulated with a SARS-CoV-2 S peptide pool (15-mer peptides with 11 amino acids overlap covering the full-length S, total 315 peptides, JPT Peptide Technologies GmbH, Berlin, Germany) at 1 µg/ml/peptide. The same volume of 40% DMSO (Sigma-Aldrich), the solution to dissolve the peptide pool, was used as the negative control. PMA/Ionomycin (Sigma-Aldrich) was used as the positive control. After overnight incubation, the cells were washed away, and biotinylated anti-mouse IFN-γ (BD) was added, and the plates were incubated for 2 hours. After washing with PBS/0.01% Tween 20, Streptavidin-HRP enzyme conjugate (Thermo Fisher Scientific) was added into the plates, which was incubated for 1 hour. After washing with PBS/0.01% Tween 20, TMB ELISPOT substrate (Mabtech, Cincinnati, OH) was added into the plates, and the spots were developed and read with an ImmunoSpot® Analyzer (Cellular Technology Limited, Cleveland, OH). The number of the S-specific spots was acquired by subtracting the number of the spots of the DMSO control wells from the number of the spots of the corresponding S peptide pool stimulation wells.

For IFN-γ and IL-4 ELISPOT of BALB/c mice, similar procedures as described above were performed using ImmunoSpot® Mouse IFN-γ and IL-4 ELISPOT kit (CTL, Shaker Heights, OH), with modifications according to the manufacturer’s protocol. Splenocytes were stimulated with two subpools (158 and 157 peptides, respectively) of the JPT’s SARS-CoV-2 S peptide pool separately. The number of the S-specific spots was acquired by adding up the numbers of the spots in each subpool-stimulated splenocytes minus the number of the spots of the DMSO control wells.

### T cell intracellular cytokine staining

Mouse splenocytes were cultured in RPMI-10 and stimulated with the S peptide pool at 1 µg/ml/peptide in the presence of GolgiStop™ and GolgiPlug™ (BD) for 6 hours. 40% DMSO and PMA/Ionomycin was used as the negative and positive control, respectively. Cells were first stained with the LIVE/DEAD™ Fixable Violet Dead Cell Stain, blocked the FcR with the TruStain FcX (Biolegend, San Diego, CA) and then stained with the fluorochrome-labeled anti-mouse CD3/CD4/CD8/CD44/CD62L mAbs (all purchased from Biolegend except CD44 from BD). Cells were then treated with Cytofix/Cytoperm (BD) and stained with fluorochrome-labeled anti-mouse IFN-γ/TNF-α/IL-2/IL-4/IL-5 mAbs (Biolegend). LSRFortessa was used to acquire the flow cytometry data, which were then analyzed with FlowJo. Percentage of cytokine^+^ T cells was calculated by subtracting the percentage of the DMSO control cells from the percentage of the corresponding S peptide pool stimulation cells.

### Multiplex immunoassay

Supernatant from the mouse splenocytes stimulated with the S peptide pool at 1 µg/ml/peptide for 24 hours was collected and cytokines were detected using a multiplex capture sandwich immunoassay. Bio-Plex Pro™ mouse cytokine Th1/Th2 assay kit (Bio-Rad, Mississauga, ON, Canada) was used. Both standards and samples were prepared following manufacturer’s instructions. The assay plate was read in a Bio-Plex MAGPIX system (Bio-Rad) and data were analyzed using the Bio-Plex Manager Software (Bio-Rad).

### SARS-CoV-2 mouse challenge

Mice were anesthetized with isofluorane and intranasally transduced with 10^11^ genomic copies of the AAV6-hACE2 or in some experiments AAV6-luciferase as control (kindly provided by Dr. S. K. Wootton, Department of Pathobiology, University of Guelph, Guelph, ON, Canada). Nine days later, the mice were anesthetized with isofluorane and intranasally challenged with 10^5^ TCID_50_ SARS-CoV-2 (SARS-CoV-2, isolate Canada/ON/VIDO-01/2020, GISAID accession number: EPI_ISL_425177). On 4dpi, mice were humanly euthanized and blood and lungs were collected. For each mouse, the left lung was sent for pathology examination, and the right lung was homogenized in DMEM-2, using a Bead Mill Homogenizer (OMNI International, Kennesaw GA). Lung homogenates were then clarified by centrifugation at 10,000 rpm for 5 minutes.

### Hamster vaccination and SARS-CoV-2 challenge

Male Syrian hamsters, aged 6 to 10 weeks, were obtained from Charles River Canada (Saint-Constant, QC, Canada). The animals were kept in Biosafety Level-2 housing until virus challenge in Biosafety Level-3 *in vivo* facility. A total of 2 groups of 16 animals (n=8/group) were immunized twice with a 3-week interval with 20µg of PTX-COVID19-B in 100 µl via intramuscular route into rear limbs (50 µl/limb). Mock animal group received an equivalent volume of phosphate-buffered saline (PBS). Three weeks post boost, animals were intranasally challenged with SARS-CoV-2 (1 x 10^4^ TCID_50_ in 100 µl per animal) under inhalated isofluorane anesthesia. Animals were monitored daily for clinical signs of disease and phenotype parameters such as weight loss and body temperature was recorded every second day. No death was recorded after the viral infection.

Four animals in both challenged group were humanly euthanized at 4 and 8 dpi for virological and histopathological analyses. Blood and major organ tissues were collected, and the tissues were separated into 2 parts, one part immediately fixed in 10% formalin, and the other part immediately frozen at −80°C until further use. The frozen tissue samples were homogenized in 1 ml DMEM-2 manually in a disposable 15 ml closed Tissue Grinder System (Thermo Fisher Scientific). 140 µl out of 1ml samples were used for RNA extraction while 500 µl of homogenates were used for quantification of SARS-CoV-2. For oral swabs, anesthetized animals were swabbed (9-11 seconds swabbing) and swabs were then introduced into 1ml DMEM-2. All oral swab samples were frozen until further processing. 500 µl of each oral swab sample was used for quantification of SARS-CoV-2.

### Determination of infectious SARS-CoV-2 titer

VeroE6 cells cultured in DMEM-10 were seeded into 96-well plates and incubated overnight at 37°C. On the following day culture medium was removed and tissue samples 10-fold serially diluted in DMEM supplemented with 1% FBS were added onto the cells. The plates were then incubated at 37°C for 1hour. After incubation lung homogenates were replaced with 100µl/well DMEM-2, and the cells were incubated at 37°C for 5 days. Cytopathic effect (CPE) was checked on day 3 and day 5. TCID_50_ was defined as the highest dilution factor of the inoculum that yielded 50% of the cells with CPE and determined by using the Spearman-Karber TCID_50_ method.

### Real-time RT-PCR

Real-time RT-PCR to quantify the genomic copies of SARS-CoV-2 in tissue homogenates was done according to the published protocol(*40*). Briefly, RNA was isolated from the tissue homogenates using QIAamp viral RNA kit (QIAGEN, Toronto, ON, Canada). Luna Universal Probe One-step RT-qPCR kit (New England Biolabs, Ipswich, MA) was used to amplify the envelope (E) gene using the following primers and probes: forward primer: ACAGGTACGTTAATAGTTAATAGCGT, reverse primer: ATATTGCAGCAGTACGCACACA, and probe CAL Fluor Orange 560-ACACTAGCCATCCTTACTGCGCTTCG-BHQ-1. The cycling conditions were 1 cycle at 60°C for 10 minutes, then 95°C for 2 minutes, followed by 44 cycles at 95°C for 10 seconds and 60°C for 15 seconds. An E gene DNA standard (pUC57-2019-nCoV-PC:E, GenScript, Piscataway, NJ) was also run at the same time for conversion of Ct value to genomic copies, by using the Rotor-Gene Q software (QIAGEN).

### Pathology

The formalin-fixed lung tissue was processed for paraffin embedding, microtomy and then stained with hematoxylin and eosin. The blocks were examined at 3 separate levels (3 separate slides). Histological sections were examined blind to vaccination status. Semiquantitative grading of lung was conducted according to Table S1.

### Statistical analysis

One-way ANOVA (Kruskal-Wallis test) followed by Dunn’s multiple comparison, two-way ANOVA followed by Sidak’s multiple comparison, two-tailed paired t test, or two-tailed unpaired t test (Mann-Whitney) were used for comparison between groups, as indicated in the figure legends. Spearman correlation test was used for correlation analysis. Logistic regression was used for determining nAb ID_50_ threshold titer that would confer 95% predicted probability of protection from productive SARS-CoV-2 infection in mice. All statistical analysis was performed by using GraphPad Prism 8. *P*<0.05 was regarded as statistically significant.

## Supplementary Materials

**Fig. S1. Expression of the SARS-CoV-2 mRNA vaccine candidates in *vitro*.**

**Fig. S2. Flow cytometry analysis of the mouse cellular immune response elicited by PTX-COVID19-B.**

**Fig. S3. AAV6-hACE2 mouse model.**

**Fig. S4. Correlation of protection afforded by PTX-COVID19-B with titers of neutralizing antibody against SARS-CoV-2 authentic virus.**

**Table S1. Lung pathology semiquantitative grading criteria.**

## Acknowledgements

We thank Drs. J.D. Bloom, J. R. Carlyle, J.E. Crowe Jr., J. M. Rini, and S. K. Wootton for providing reagents, Drs. S. Gray-Owen and N. Christie for their help in BSL-3 work, Dr. J. LaPierre, Mses. T. McCook, J. Kontogiannis, V. Chan, S. Johnson, L. Kent, J. Suarez, Mrs. F. Giuliano and J. Reid for mouse maintenance, vaccination and sampling, Mses. A. Fong and A. Antenucci for their help in pathology, Ms. M. Sharma for her help in multiplex immunoassay and Dr. B.H. Barber for helpful discussions. We thank and acknowledge the full team at Providence Therapeutics for its work on developing this vaccine and the team at Genevant Sciences for formulation of the vaccines used in these animal studies.

## Funding

This work was funded by the National Research Council of Canada–Industrial Research Assistance Program (NRC-IRAP) to Providence Therapeutics Holdings, Inc.

Canadian Institutes of Health Research (MAO)

Ontario HIV Treatment Network (MAO)

Li Ka Shing Knowledge Institute (MAO)

Juan and Stefania fund for COVID-19 and other virus infections (MAO)

Canadian Institutes of Health Research VRI-172711 (ACG)

Canadian Institutes of Health Research, Canada Research Chair, Tier 1, in Functional Proteomics (ACG)

Krembil Foundation and Royal Bank of Canada to the Sinai Health System Foundation (ACG)

Canadian Institutes of Health Research (KM)

Canadian Institutes of Health Research (AB)

Natural Sciences and Engineering Research Council of Canada (AB)

## Author contributions

Conceptualization: JL, JAA, JAS, NMO, EGM, MAO

Methodology: JL, PB, RS, BG, GB, KC, JAA, JAS, AB, KM, SM, RAK, MSP, NMO, A-CG, EG, MAO

Investigation: JL, PB, RS, BG, GB, BR, JAA, RL, LY, SKA, SC, MN, YA, QH, MSP

Visualization: JL, PB, RS, KC, JAS, YA, AH, MSP, NMO

Funding acquisition: SM, RAK, NMO, A-CG, EGM, MAO

Supervision: SM, RAK, NMO, A-CG, EGM, MAO

Writing – original draft: JL

Writing – review & editing: All authors

## Data and materials availability

All data are available in the main text or the supplementary material. Materials are available under a material transfer agreement.

## References and Notes

1. E. Dong, H. Du, L. Gardner, An interactive web-based dashboard to track COVID-19 in real time. The Lancet. Infectious diseases 20, 533–534 (2020).

2. J. H. Beigel et al., Remdesivir for the Treatment of Covid-19-Final Report. The New England journal of medicine 383, 1813–1826 (2020).

3. P. Chen et al., SARS-CoV-2 Neutralizing Antibody LY-CoV555 in Outpatients with Covid-19. The New England journal of medicine 384, 229–237 (2021).

4. P. Horby et al., Dexamethasone in Hospitalized Patients with Covid-19. The New England journal of medicine 384, 693–704 (2021).

5. M. J. Joyner et al., Convalescent Plasma Antibody Levels and the Risk of Death from Covid-19. The New England journal of medicine, (2021).

6. D. M. Weinreich et al., REGN-COV2, a Neutralizing Antibody Cocktail, in Outpatients with Covid-19. The New England journal of medicine 384, 238–251 (2021).

7. L. S. o. H. T. Medicine.

8. W. H. Organization. (2021), vol. 2021.

9. Y. Liu et al., Neutralizing Activity of BNT162b2-Elicited Serum - Preliminary Report. The New England journal of medicine, (2021).

10. T. Tada, et al., Neutralization of viruses with European, South African, and United States SARS-CoV-2 variant spike proteins by convalescent sera and BNT162b2 mRNA vaccine-elicited antibodies. bioRxiv : the preprint server for biology, (2021).

11. Z. Wang et al., mRNA vaccine-elicited antibodies to SARS-CoV-2 and circulating variants. Nature, (2021).

12. K. Wu et al., Serum Neutralizing Activity Elicited by mRNA-1273 Vaccine - Preliminary Report. The New England journal of medicine, (2021).

13. L. Corey, J. R. Mascola, A. S. Fauci, F. S. Collins, A strategic approach to COVID-19 vaccine R&D. Science (New York, N.Y.) 368, 948–950 (2020).

14. B. Coutard et al., The spike glycoprotein of the new coronavirus 2019-nCoV contains a furin-like cleavage site absent in CoV of the same clade. Antiviral research 176, 104742 (2020).

15. M. Hoffmann et al., SARS-CoV-2 Cell Entry Depends on ACE2 and TMPRSS2 and Is Blocked by a Clinically Proven Protease Inhibitor. Cell 181, 271–280.e278 (2020).

16. A. C. Walls et al., Structure, Function, and Antigenicity of the SARS-CoV-2 Spike Glycoprotein. Cell 181, 281–292.e286 (2020).

17. D. Wrapp, et al., Cryo-EM structure of the 2019-nCoV spike in the prefusion conformation. Science (New York, N.Y.) 367, 1260–1263 (2020).

18. F. Wu et al., A new coronavirus associated with human respiratory disease in China. Nature 579, 265–269 (2020).

19. P. Zhou et al., A pneumonia outbreak associated with a new coronavirus of probable bat origin. Nature 579, 270–273 (2020).

20. M. Hoffmann, H. Kleine-Weber, S. Pöhlmann, A Multibasic Cleavage Site in the Spike Protein of SARS-CoV-2 Is Essential for Infection of Human Lung Cells. Molecular cell 78, 779–784.e775 (2020).

21. T. P. Peacock, et al., The furin cleavage site of SARS-CoV-2 spike protein is a key determinant for transmission due to enhanced replication in airway cells. bioRxiv : the preprint server for biology, (2020).

22. B. A. Johnson et al., Loss of furin cleavage site attenuates SARS-CoV-2 pathogenesis. Nature, (2021).

23. C. O. Barnes et al., SARS-CoV-2 neutralizing antibody structures inform therapeutic strategies. Nature 588, 682–687 (2020).

24. P. J. M. Brouwer et al., Potent neutralizing antibodies from COVID-19 patients define multiple targets of vulnerability. Science (New York, N.Y.) 369, 643–650 (2020).

25. B. Ju et al., Human neutralizing antibodies elicited by SARS-CoV-2 infection. Nature 584, 115–119 (2020).

26. L. Liu et al., Potent neutralizing antibodies against multiple epitopes on SARS-CoV-2 spike. Nature 584, 450–456 (2020).

27. D. F. Robbiani et al., Convergent antibody responses to SARS-CoV-2 in convalescent individuals. Nature 584, 437–442 (2020).

28. T. F. Rogers et al., Isolation of potent SARS-CoV-2 neutralizing antibodies and protection from disease in a small animal model. Science (New York, N.Y.) 369, 956–963 (2020).

29. R. Shi et al., A human neutralizing antibody targets the receptor-binding site of SARS-CoV-2. Nature 584, 120–124 (2020).

30. S. J. Zost et al., Potently neutralizing and protective human antibodies against SARS-CoV-2. Nature 584, 443–449 (2020).

31. J. Gergen, B. Petsch, mRNA-Based Vaccines and Mode of Action. Current topics in microbiology and immunology, (2021).

32. K. S. Corbett et al., SARS-CoV-2 mRNA vaccine design enabled by prototype pathogen preparedness. Nature 586, 567–571 (2020).

33. L. R. Baden et al., Efficacy and Safety of the mRNA-1273 SARS-CoV-2 Vaccine. The New England journal of medicine 384, 403–416 (2021).

34. F. P. Polack et al., Safety and Efficacy of the BNT162b2 mRNA Covid-19 Vaccine. The New England journal of medicine 383, 2603–2615 (2020).

35. B. Korber et al., Tracking Changes in SARS-CoV-2 Spike: Evidence that D614G Increases Infectivity of the COVID-19 Virus. Cell 182, 812–827.e819 (2020).

36. A. C. Walls et al., Cryo-electron microscopy structure of a coronavirus spike glycoprotein trimer. Nature 531, 114–117 (2016).

37. M. A. Tortorici et al., Structural basis for human coronavirus attachment to sialic acid receptors. Nature structural & molecular biology 26, 481–489 (2019).

38. A. G. Wrobel et al., SARS-CoV-2 and bat RaTG13 spike glycoprotein structures inform on virus evolution and furin-cleavage effects. Nature structural & molecular biology 27, 763–767 (2020).

39. X. Yang et al., Modifications that stabilize human immunodeficiency virus envelope glycoprotein trimers in solution. Journal of virology 74, 4746–4754 (2000).

40. A. Banerjee et al., Isolation, Sequence, Infectivity, and Replication Kinetics of Severe Acute Respiratory Syndrome Coronavirus 2. Emerging infectious diseases 26, 2054–2063 (2020).

41. P. H. England. (2020), vol. 2021.

42. H. Tegally et al., Detection of a SARS-CoV-2 variant of concern in South Africa. Nature 592, 438–443 (2021).

43. N. R. Faria et al., Genomics and epidemiology of the P.1 SARS-CoV-2 lineage in Manaus, Brazil. Science (New York, N.Y.), (2021).

44. J. R. Habel et al., Suboptimal SARS-CoV-2-specific CD8(+) T cell response associated with the prominent HLA-A*02:01 phenotype. Proceedings of the National Academy of Sciences of the United States of America 117, 24384–24391 (2020).

45. J. C. Law et al., Systematic Examination of Antigen-Specific Recall T Cell Responses to SARS-CoV-2 versus Influenza Virus Reveals a Distinct Inflammatory Profile. Journal of immunology (Baltimore, Md. :1950) 206, 37–50 (2021).

46. G. Breton et al., Persistent cellular immunity to SARS-CoV-2 infection. The Journal of experimental medicine 218, (2021).

47. A. T. Tan et al., Early induction of functional SARS-CoV-2-specific T cells associates with rapid viral clearance and mild disease in COVID-19 patients. Cell reports 34, 108728 (2021).

48. A. Bonifacius et al., COVID-19 immune signatures reveal stable antiviral T cell function despite declining humoral responses. Immunity 54, 340–354.e346 (2021).

49. B. Israelow et al., Mouse model of SARS-CoV-2 reveals inflammatory role of type I interferon signaling. The Journal of experimental medicine 217, (2020).

50. J. F. Chan, et al., Simulation of the Clinical and Pathological Manifestations of Coronavirus Disease 2019 (COVID-19) in a Golden Syrian Hamster Model: Implications for Disease Pathogenesis and Transmissibility. Clinical infectious diseases : an official publication of the Infectious Diseases Society of America 71, 2428–2446 (2020).

51. S. F. Sia et al., Pathogenesis and transmission of SARS-CoV-2 in golden hamsters. Nature 583, 834–838 (2020).

52. J. Draize, G. Woodard, H. Calevery, Methods for the study of irritation and toxicity of substances applied topically to the skin and mucous membranes. Journal of pharmacology and Experimental therapeutics 82, 14 (1944).

53. A. B. Vogel et al., BNT162b vaccines protect rhesus macaques from SARS-CoV-2. Nature, (2021).

54. F. Krammer, SARS-CoV-2 vaccines in development. Nature 586, 516–527 (2020).

55. K. Subbarao, B. R. Murphy, A. S. Fauci, Development of effective vaccines against pandemic influenza. Immunity 24, 5–9 (2006).

56. K. S. Corbett et al., Evaluation of the mRNA-1273 Vaccine against SARS-CoV-2 in Nonhuman Primates. The New England journal of medicine 383, 1544–1555 (2020).

57. M. Guebre-Xabier et al., NVX-CoV2373 vaccine protects cynomolgus macaque upper and lower airways against SARS-CoV-2 challenge. Vaccine 38, 7892–7896 (2020).

58. A. Muik et al., Neutralization of SARS-CoV-2 lineage B.1.1.7 pseudovirus by BNT162b2 vaccine-elicited human sera. Science (New York, N.Y.), (2021).

59. C. Keech et al., Phase 1-2 Trial of a SARS-CoV-2 Recombinant Spike Protein Nanoparticle Vaccine. The New England journal of medicine 383, 2320–2332 (2020).

60. J. Sadoff et al., Interim Results of a Phase 1-2a Trial of Ad26.COV2.S Covid-19 Vaccine. The New England journal of medicine, (2021).

61. J. H. Lee, G. Ozorowski, A. B. Ward, Cryo-EM structure of a native, fully glycosylated, cleaved HIV-1 envelope trimer. Science (New York, N.Y.) 351, 1043–1048 (2016).

62. D. M. McCraw et al., Structural analysis of influenza vaccine virus-like particles reveals a multicomponent organization. Scientific reports 8, 10342 (2018).

63. M. S. A. Gilman et al., Transient opening of trimeric prefusion RSV F proteins. Nature communications 10, 2105 (2019).

64. Y. Watanabe, et al., Native-like SARS-CoV-2 spike glycoprotein expressed by ChAdOx1 nCoV-19/AZD1222 vaccine. bioRxiv : the preprint server for biology, (2021).

65. A. C. Walls et al., Elicitation of Potent Neutralizing Antibody Responses by Designed Protein Nanoparticle Vaccines for SARS-CoV-2. Cell 183, 1367–1382.e1317 (2020).

66. H. X. Tan et al., Immunogenicity of prime-boost protein subunit vaccine strategies against SARS-CoV-2 in mice and macaques. Nature communications 12, 1403 (2021).

67. C. Rydyznski Moderbacher et al., Antigen-Specific Adaptive Immunity to SARS-CoV-2 in Acute COVID-19 and Associations with Age and Disease Severity. Cell 183, 996–1012.e1019 (2020).

68. K. McMahan et al., Correlates of protection against SARS-CoV-2 in rhesus macaques. Nature 590, 630–634 (2021).

69. S. Bedoui, W. R. Heath, S. N. Mueller, CD4(+) T-cell help amplifies innate signals for primary CD8(+) T-cell immunity. Immunological reviews 272, 52–64 (2016).

70. A. Ciurea, L. Hunziker, P. Klenerman, H. Hengartner, R. M. Zinkernagel, Impairment of CD4(+) T cell responses during chronic virus infection prevents neutralizing antibody responses against virus escape mutants. The Journal of experimental medicine 193, 297–305 (2001).

71. S. Crotty, T Follicular Helper Cell Biology: A Decade of Discovery and Diseases. Immunity 50, 1132–1148 (2019).

72. A. D. Redd, et al., CD8+ T cell responses in COVID-19 convalescent individuals target conserved epitopes from multiple prominent SARS-CoV-2 circulating variants. medRxiv : the preprint server for health sciences, (2021).

73. T. J. Ruckwardt, K. M. Morabito, B. S. Graham, Immunological Lessons from Respiratory Syncytial Virus Vaccine Development. Immunity 51, 429–442 (2019).

74. K. T. Abe et al., A simple protein-based surrogate neutralization assay for SARS-CoV-2. JCI insight 5, (2020).

75. M. Stuible et al., Rapid, high-yield production of full-length SARS-CoV-2 spike ectodomain by transient gene expression in CHO cells. Journal of biotechnology 326, 21–27 (2021).

