## Supplementary materials for "Preclinical evaluation of a SARS-CoV-2 mRNA vaccine PTX-COVID19-B"

**Supplementary Figures:**

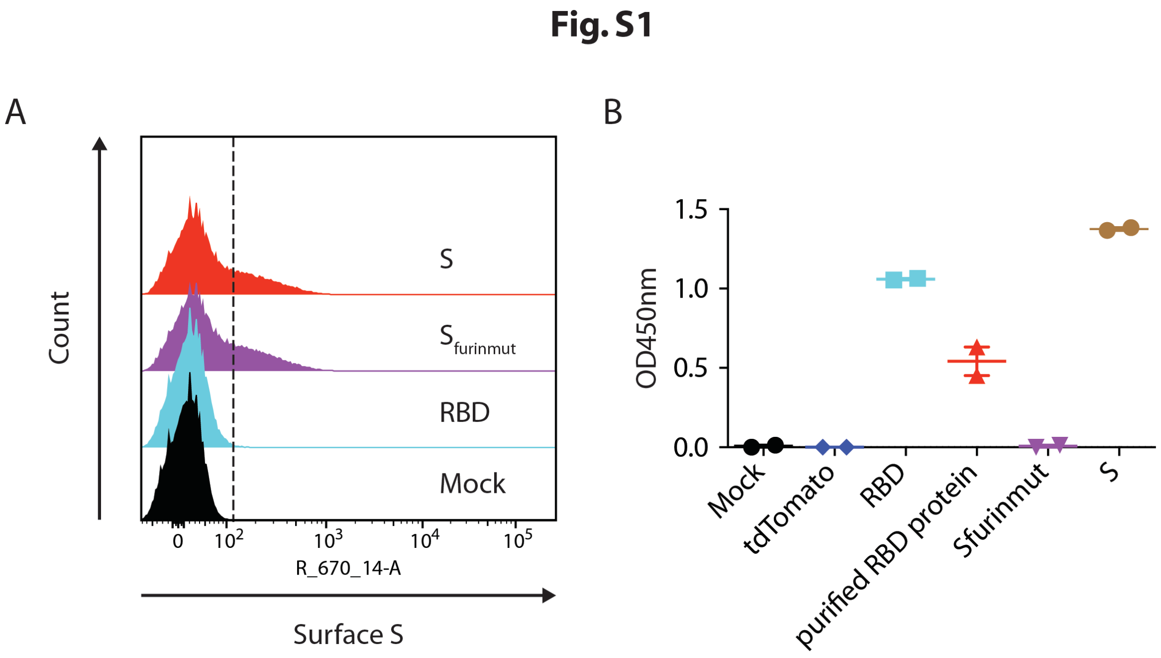


**Figure S1. Expression of the SARS-CoV-2 mRNA vaccine candidates in *vitro*.** HEK293T cells were transfected with the mRNA vaccine constructs and the expression of these constructs on the cell surface was detected by flow cytometry in **(A)** and in the supernatant by ELISA in **(B)** by using an RBD-specific neutralizing mAb COV2-2165 as the detection antibody**.** Shown in (A) is a flow cytometry histogram indicating the expression of the constructs on the surface of the transfected HEK293T cells. The dashed line is the gating boundary between $S^+$ and $S^-$ cells. The result is a representative of 3 independent experiments. Shown in (B) are $OD_{450}$ readings. Mock: mock transfection. Purified RBD protein: 100ng/ml of purified RBD protein. For each group, one symbol represents one of 2 technical replicates, long line indicates the mean and the error bars indicate SEM.

**Fig. S2**

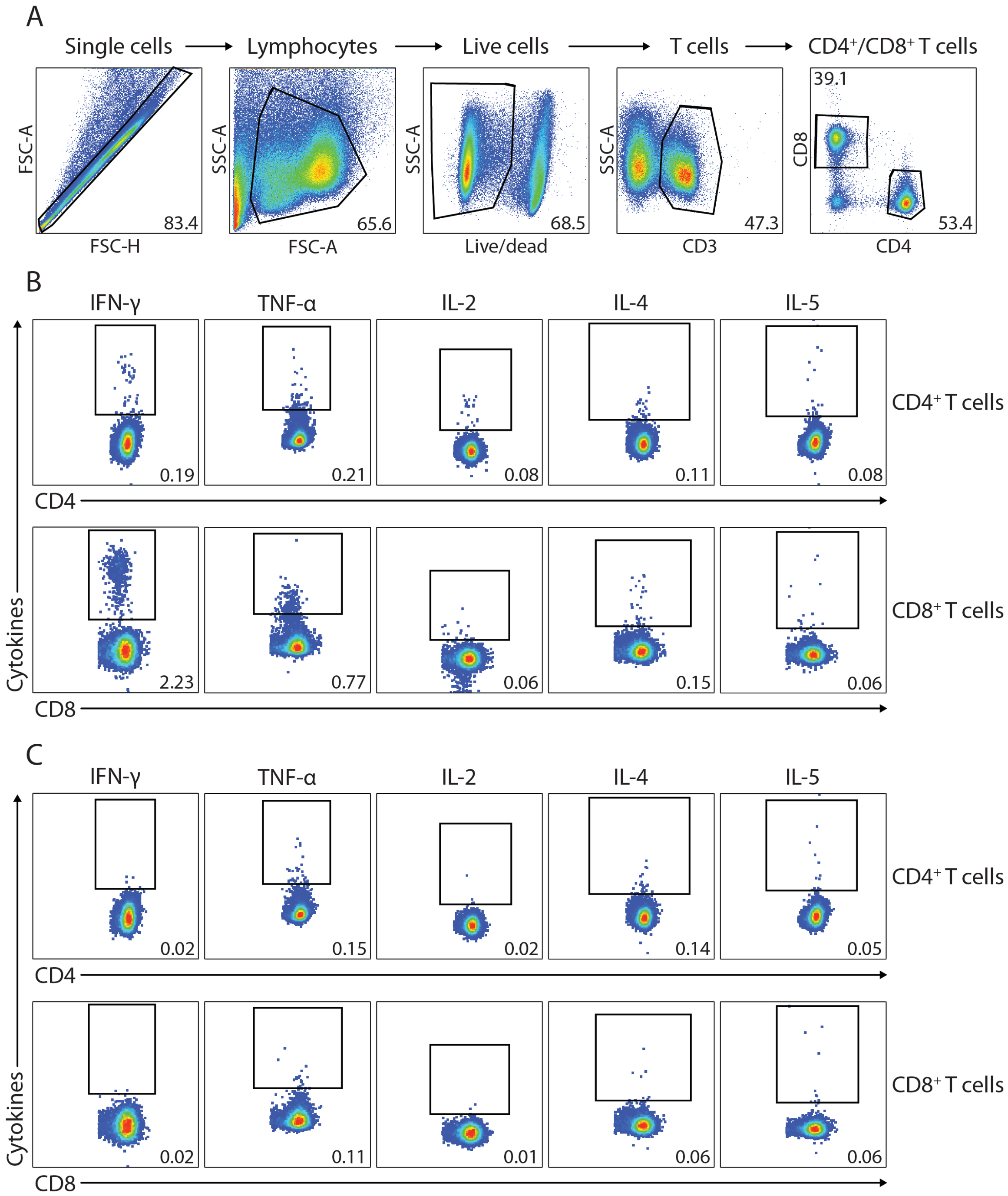


**Figure S2. Flow cytometry analysis of the mouse cellular immune response elicited by PTX-COVID19-B.** Splenocytes from vaccinated mice in Fig. 4 were stimulated with a SARS-CoV-2 S peptide pool for 6 hours in the presence of GolgiPlug and GolgiStop, stained with mAbs and analyzed with flow cytometry. **(A)** Gating strategy for intracellular cytokine staining. **(B)** and **(C)** Representative flow cytometry graphs of cytokine production of T cells from a PTX-COVID19-B vaccinated mouse in (B) and a control mouse receiving tdTomato mRNA in (C). The number in each graph is the percentage of gated cells in its parental cell population, e.g. percentage of cytokine-producing $CD4^+$ or $CD8^+$ T cells in total $CD4^+$ or $CD8^+$ T cells.

**Fig. S3**

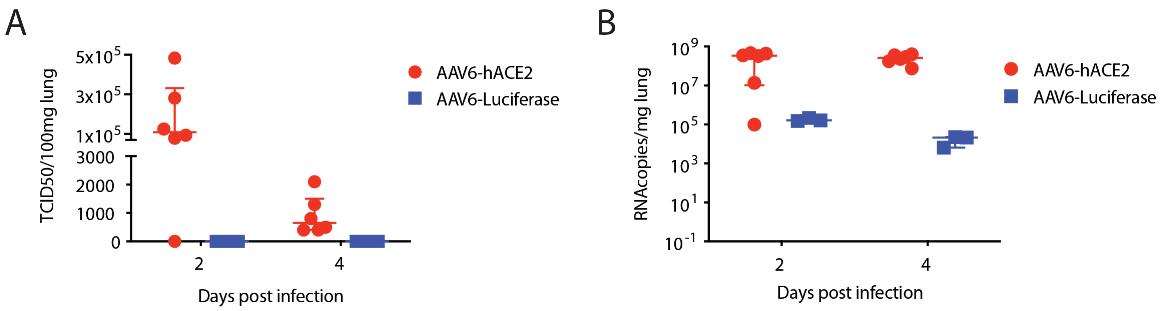


**Figure S3. AAV6-hACE2 mouse model.** Female C57BL/6 mice were intranasally transduced with AAV6-hACE2 or AAV6-Luciferase. Nine days after the transduction, mice were intranasally challenged with SARS-CoV-2. Two and 4 days after the challenge, mice were humanly euthanized and lungs were collected from the mice. **(A)** Amount of infectious SARS-CoV-2 and **(B)** SARS-CoV-2 RNA in the lungs of the mice transduced with AAV6-hACE2 or AAV6-Luciferase. Shown in (A) are $TCID_{50}$/100mg lung tissue and (B) RNA copies/mg lung tissue. N=6 for AAV6-hACE2 group per time point, n=3 for AAV6-Luciferase group per time point. Each symbol represents one mouse. For each group, the long horizontal lines indicates the medians and the short lines below and above the median indicate the 25% and 75% percentile.

### Fig. S4

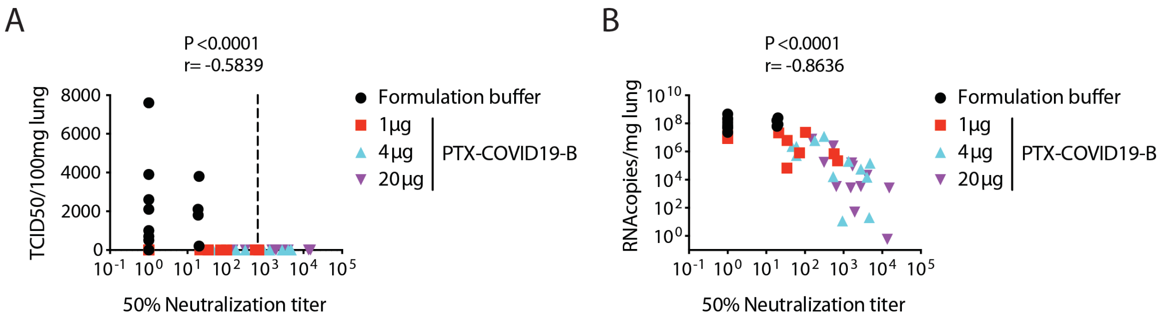


**Figure S4. Correlation of protection afforded by PTX-COVID19-B with titers of neutralizing antibody against SARS-CoV-2 authentic virus.** Data from vaccinated and SARS-CoV-2 challenged mice in Fig. 5 were analyzed. Shown is correlation of 50% neutralization titers ($ID_{50}$) of the sera collected at 4 days after SARS-CoV-2 challenge with **(A)** the amount of infectious SARS-CoV-2 virus and **(B)** SARS-CoV-2 RNA in the lungs of the mice. *P* and r values are determined by Spearman correlation test. Dashed line in (A) indicates the threshold $ID_{50}$ titer that predicts 95% probability of protection from productive SARS-CoV-2 infection as determined by logistic regression.

**Table S1. Lung pathology semiquantitative grading criteria**

| Grade | Inflammation | Histologic findings |
|---|---|---|
| 0 | None | No inflammation |
| 1 | Borderline | Indistinct or slight inflammatory cell infiltrates |
| 2 | Mild, or airway predominant inflammation | Noncircumferential perivascular/peribronchiolar inflammatory infiltrates, mostly sparse; inflammatory changes in respiratory mucosa of parenchymal airways. This grade also represents marked bronchitis without substantial parenchyma involvement |
| 3 | Moderate | Moderate and multifocal pneumonitis; inflammatory changes in respiratory mucosa of parenchymal airways |
| 4 | Severe | Marked and extensive pneumonitis; inflammatory changes in respiratory mucosa of parenchymal airways |
